# Alpha-range neural oscillations are associated with multiple dynamic brain networks

**DOI:** 10.64898/2026.10.12.758569

**Authors:** J.S. Nomi, T.V. Román-López, F.Y. Chang, A. Friedman-Jha, H.V. Truong, J.P. Diaz-Fong, C. Chang, S.L. Karalunas, S. Phadikar, V.D. Calhoun, A. Dillon, S.K. Loo, A. Lenartowicz

## Abstract

We asked whether alpha range (8-12 Hz) EEG neural oscillations, a robust indicator of cortical excitability, are associated with dynamic BOLD brain networks. Two independent EEG-fMRI resting-state datasets (discovery: *n* = 94 and replication: *n* = 33) show that low-and high-alpha periods are associated with dynamic BOLD coactivation patterns (dCAPS) involving multiple dynamic networks rather than a single static network. Unique BOLD dCAPs consisting of visual cortex activation along with sensorimotor and attention networks in addition to the posterior thalamus, were mainly associated with periods of low-alpha power. However, overlapping BOLD dCAPs consisting of default, control, and attention networks were associated with periods of both low-and high-alpha power. The results suggest low-alpha dCAPS contain unique temporal-dependent configurations related to visual, sensorimotor, and attention networks while both low-and high-alpha dCAPs consist of overlapping temporal-dependent configurations involving CON, DMN, and attention networks involved in internal and external cognition. These findings reconcile discrepancies across static connectivity EEG-fMRI findings by showing that distinct transient BOLD configurations are associated with similar patterns of alpha EEG modulation in visual cortex, providing a dynamic framework for the gating function of alpha oscillations.

## Introduction

Alpha range (8-12Hz) neural oscillations are a robust indicator of cortical excitability^1–4^, associated with a gating function critical to shaping neural communication within cortex^5–7^. For instance, in the context of selective attention, decreases in alpha power accompany activation of regions involved in target processing, whereas increases in alpha power can be observed in cortex associated with distractor processing along with a lower-excitability state^3,8^. The relationship between alpha oscillations and blood-oxygenation level-dependent (BOLD) functional magnetic resonance imaging (fMRI) activity is less clear. One possibility is that alpha power may vary with BOLD activity related to attention systems such as the dorsal or ventral attention networks (DAN, VAN)^9^. Indeed, decreases of alpha power in several concurrent EEG-fMRI studies have been associated with increased BOLD activation in distributed visual and fronto-parietal regions (including the DAN) at rest^10,11^ and during stimulus processing^12–14^. However, increases of alpha power, have been associated with BOLD activity within the default mode network (DMN)^14–17^, cingulo-opercular network^18,19^ (i.e., overlapping with the salience network (SN)), and thalamo-cortical circuitry^20,21^. Further, global EEG alpha fluctuations have been linked to a broad physiological arousal response, including changes in global fMRI signal^22,23^.

The variability in reported BOLD activity associated with alpha oscillations is intriguing. While it could reflect measurement error or methodological differences^24,25^, it could also indicate the dynamic nature of the brain itself, with alpha oscillations capturing temporal-and context-dependent fluctuations in cortical excitability^4,26^, that translate into context-dependent fluctuations in BOLD signal. The reported EEG-fMRI results are based largely on static functional connectivity (sFC) methods^10,27^, designed to identify the dominant association between EEG and fMRI signals, even if others are present transiently. However, the variety of BOLD activity associated with alpha oscillations suggests a dynamic, rather than static, relationship between the two measures. In other words, alpha power decreases may sometimes be associated with BOLD activity increases in visual cortex, while other times extending into, sensory motor, and fronto-parietal regions^10–14^. Whereas alpha power increases may be associated with lower visual and DMN BOLD activation^14–17^ because of a shift from external to internal processing or because of thalamo-cortical^20,21^ down regulation of visual cortex BOLD activity with a change in arousal^20,22^. Multiple EEG-fMRI associations could occur for periods of high-or low-alpha transiently within the same time series, though sFC methods would not detect such brain dynamics.

Few EEG-fMRI studies have quantified dynamic relationships between alpha oscillations and BOLD activity^28–31^. Multivariate functional connectivity analyses point to widespread network associations^29^, that vary with alpha sources^30,32^ and show transient dynamics^31^, supporting the hypothesis that multiple, transient associations between alpha power and BOLD activity exist. However, functional connectivity dynamics examines continuous relationships between alpha power with the BOLD signal. It does not differentiate between discrete periods of low-alpha power and high-alpha power that may facilitate unique or overlapping dynamic brain configurations. Dynamic coactivation pattern analysis (dCAP)^33^ is an alternate approach that has been effectively deployed to identify multiple patterns of BOLD activity associated with discrete timepoints of a thresholded signal throughout an fMRI timeseries. This generally involves clustering BOLD images associated with strong activity above a predetermined threshold related to a targeted brain area. This would be advantageous for examining BOLD dynamics associated with periods of high-alpha power. However, dCAP analysis has also been used to identify brain activity associated with the troughs or valleys of the BOLD signal^34^. This would be advantageous for examining BOLD dynamics associated with periods of low alpha-power. Results suggest that the peaks and troughs of the BOLD signal identify unique and overlapping brain configurations^34^. Applying these two contrasting dCAP approaches to concurrent EEG-fMRI data may help to reconcile the heterogenous sFC EEG-fMRI findings by quantifying if different BOLD configurations can be associated with high-and low-alpha power within the context of a single resting-state scan.

Here, we leverage dCAP analyses in two independently acquired multi-modal EEG-fMRI datasets to identify dynamic brain states associated with periods of high and low occipital alpha power. Using a discovery and replication approach, we show that BOLD-alpha power relationships are inherently dynamic, comprising multiple transient BOLD coactivation states during both high-and low-alpha periods. These states encompass previously reported sFC associations while also revealing more spatially extensive coactivation patterns spanning multiple canonical cortical networks and subcortical structures, including the anterior and posterior thalamus. Across datasets, unique dCAPs associated with low-alpha power consistently involved visuo-parietal networks, with varying coactivation of posterior thalamus and association cortices, suggesting dynamic temporal-dependent reconfiguration. Dynamic overlapping dCAPs associated with both low-and high-alpha power were spatially heterogeneous but consistent with internally/externally directed cognition. These findings demonstrate multiple manifestations of alpha BOLD activity modulation in the visual cortex. These findings also reconcile seemingly inconsistent reports of alpha-BOLD associations by showing that they may represent distinct transient brain states rather than competing functional interpretations.

## Results

### Alpha power coactivation patterns are dynamic

To identify multiple functional networks associated with alpha power during rest, we applied dCAP analysis^33^ to resting-state fMRI signals in Dataset 1 (*n* = 94) (**S Table 1**). On the EEG side, alpha power was computed from occipital electrodes, normalized, and averaged across electrodes to create a single alpha power time series. This averaged time series was then convolved with a hemodynamic response function (HRF) to align it with the timescale of the fMRI data (**Fig. 1ab**). On the fMRI side, BOLD signals were summarized using 400 ROI time series from the Schaefer 400 ROI Kong 17 Network parcellation^35^ and 54 subcortical ROI time series from the Tian subcortical parcellation^36^ (**Fig 1a**). Timepoints corresponding to the top 15% and bottom 15% of alpha power time series (for thresholds of 10% see **S Figures 1-2**; for 20% see **S Figures 3-4**) were then identified, representing periods of high-alpha (“peaks”) and low-alpha (“troughs”) states respectively (for thresholds of 10% see **S Figures 1-2**; for 20% see **S Figures 3-4**) (**Fig 1e**). The corresponding fMRI timepoints were concatenated across all individual subject datasets and dCAP states were identified by k-means clustering^37^. The procedure revealed that high-alpha and low-alpha states were each associated with five distinct clusters of BOLD spatial patterns, varying in frequency of occurrence between 16% and 24% (**Fig. 2**, right panels). We thus labeled the former as dynamic high-alpha coactivation patterns (dHAPs) and the latter as dynamic low-alpha coactivation patterns (dLAPs). To ensure that these dCAPs are not random noise, we additionally phase-scrambled the fMRI time series - preserving the mean, standard deviation, and autocorrelation - and repeated the analyses^38,39^ (see supplemental materials, **S Figures 5-6**). We also repeated the dCAP results without global signal regression (**S Figures 7-8**). The results confirmed that fluctuating resting-state alpha power is associated with dCAPs for both high-alpha and low-alpha time periods.

**Figure 1:**
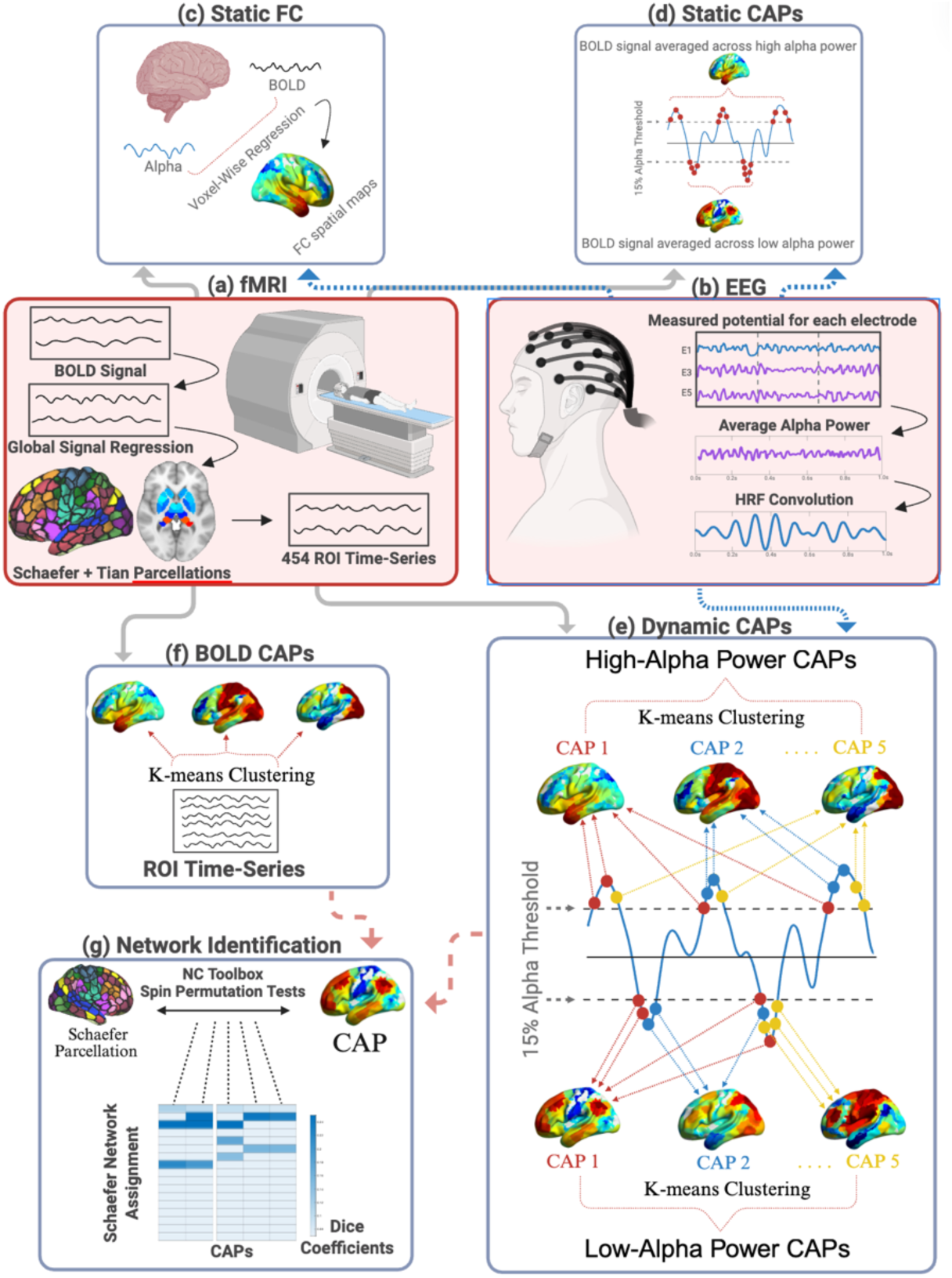
Multi-modal EEG-fMRI static and dynamic coactivation pattern analysis (Static CAPs and Dynamic CAPs) workflow. (a) fMRI. (second row, left): 454 ROI time series from the Schaefer 400 ROI Kong 17 Network parcellation^35^ and the Tian subcortical parcellation^36^ were extracted from preprocessed BOLD data. **(b) EEG** (second row, right): Average alpha power was calculated across occipital electrodes then convolved with a hemodynamic response function (HRF). **(c) Static FC** (top row, left): Voxel-wise static functional connectivity (Static FC) was calculated between the convolved alpha time series and the BOLD signal. **(d) Static CAPs** (top row, right): ROI-wise static coactivation patterns (Static CAPs) were calculated by averaging BOLD signal TRs across periods of high-and low-alpha power. **(e) Dynamic CAPs** (bottom right): TRs associated with periods of high-and low-alpha power were subjected to k-means clustering. BOLD TRs clustered across periods of high-alpha power produced High-Alpha Power dCAPs and TRs clustered across periods of low-alpha power produced low-alpha power dCAPs. **(f) BOLD CAPs** (third row, left): K-means clustering was applied to all BOLD TRs to produce CAPs that were not associated with alpha power. **(g) Network Identification** (bottom, left): The network correspondence toolbox (NCT)^40^ was used to categorize canonical BOLD network activity using spin permutation testing and dice coefficients for dynamic CAPs and BOLD CAPs.

**Figure 2:**
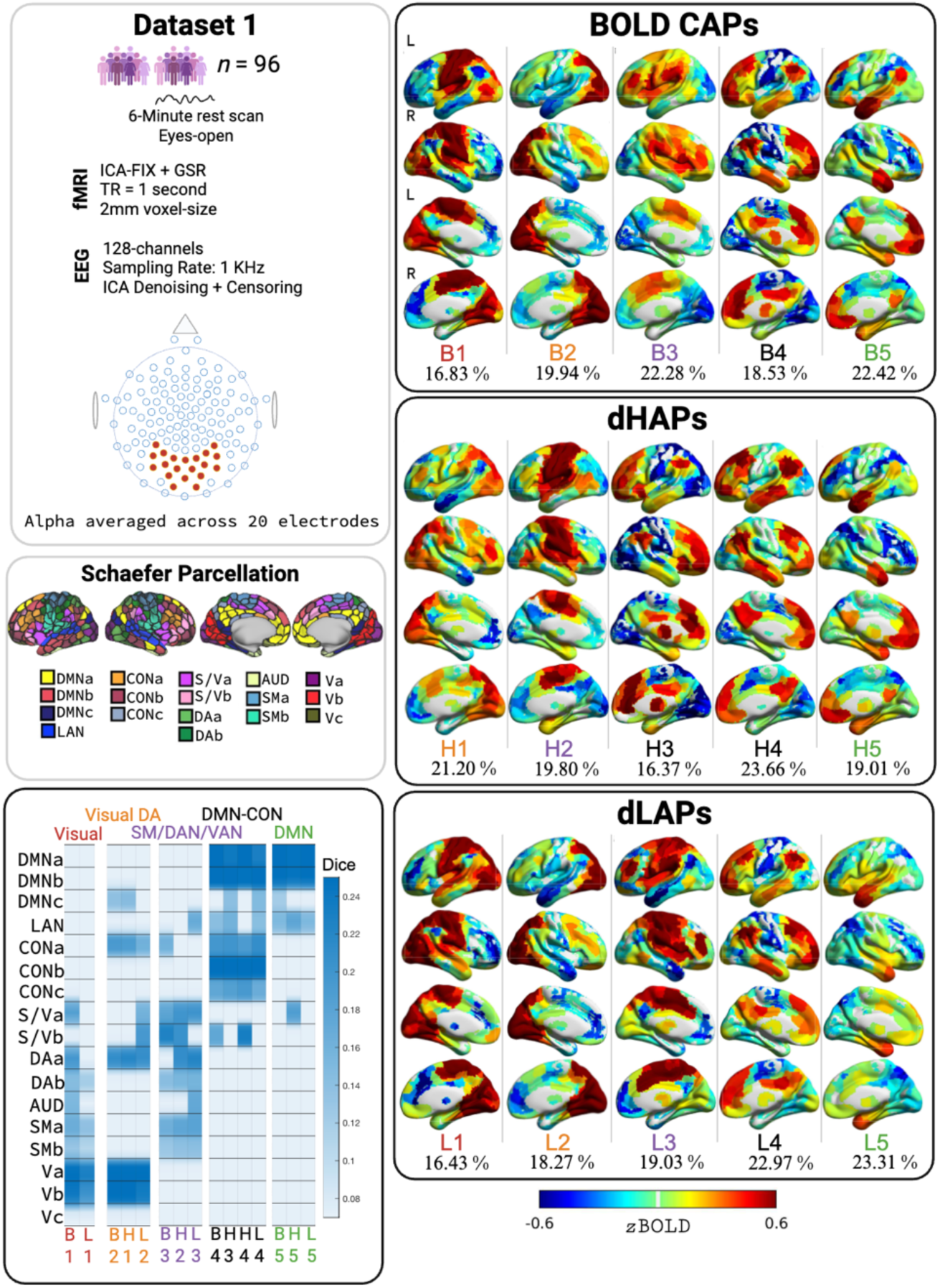
Dataset 1 results. Top Left: Acquisition, preprocessing, and demographic data for Dataset 1. **Middle Left**: Schaefer 400 ROI Kong 17 Network Parcellation^35^ was used to identify canonical BOLD network activation using spin permutation tests and Dice Coefficients. **Bottom Left**: Dice Coefficients from spin permutation tests indicating the overlap between BOLD CAPs and network assignment from the Schaefer Parcellation. Coefficients are only shown when surpassing a threshold of *p* < 0.05. **Top Right**: BOLD CAPs not associated with alpha power identified by using k-means clustering on all BOLD TRs. The frequency that each CAP occurs is shown in percentages. **Middle Right:** Dynamic high-alpha coactivation patterns (dHAPs). **Bottom Right:** Dynamic low-alpha power coactivation patterns (dLAPs). GSR = global signal regression; DMN = default mode network; LAN = language network; CON = control network; S/V = salience network/ventral attention network; DA = dorsal attention network; AUD = auditory network; SM = somatomotor network; V = visual network; a/b/c = subnetwork a, subnetwork b, subnetwork c; B1, B2, etc. = BOLD CAP1, BOLD CAP2, etc.; H1, H2, etc. = dHAP1, dHAP2, etc.; L1, L2, etc. = dLAP1, dLAP2, etc.

### High-alpha and low-alpha dCAPs differ in overlap with canonical BOLD networks

We next qualified the spatial patterns of the five-cluster solutions for dLAPs and dHAPs. We assessed the spatial similarity of each dCAP to the Schaefer 400 ROI Kong 17 canonical BOLD network parcellation^35^ via dice coefficients and spin test permutations (Network Correspondence Toolbox (NCT)^40^, **Fig. 1g**). This revealed clear overlap across dLAPs and dHAPs (**Fig 2**, heatmaps, bottom-left), raising the possibility of non-uniqueness between dCAPs. This may be due to temporal overlap introduced between high-alpha and low-alpha, or intervening timepoints, when convolving with the HRF. For an EEG-independent reference solution, we repeated the dCAP analysis on all timepoints of the fMRI BOLD signal, irrespective of alpha state, with the goal of identifying the dominant BOLD CAP states present during the recordings (**Fig. 1f**). This EEG-independent reference solution represents a clustering of all time points^34^, in contrast to dLAPs and dHAPs. These EEG-independent clusters were then used as a reference to identify unique and shared dCAPs across high-and low-alpha states.

The EEG-independent solution (**Fig. 2**, top-right) revealed five dominant BOLD CAPs (B1:B5, see **S Figure 9** for phase-scrambled results and see **S Figure 10** for results without global signal regression) differing in canonical network overlap (**Fig. 2**, heatmaps, bottom-left). These CAPs included two visual system configurations: (B1) coactivation of visual networks with auditory, SM and attention (DAN/VAN) networks (labeled “visual”, frequency of occurrence (i.e., how often a CAP appeared) = 17%), and (B2) coactivation of visual networks with DAN, control and DMN (“visual-DAN”, 20%). An additional CAP (B3) was characterized by a complementary pattern of SM, DAN and VAN activation but without visual networks (“SM/DAN/VAN”, 22%). The remaining two CAPs involved DMN activation: (B4) DMN with control network coactivation (“DMN-CON”, 18%) and (B5) DMN without control network activation (instead including the SN and language network) (“DMN”, 22%). A chi-squared test showed that there was no difference in the frequency of occurrence between dHAPs, dLAPs, and BOLD CAPs for Dataset 1 (χ²(8, *n* = 298) = 2.97, *p* = 0.94, Cramér’s V = 0.07).

These five BOLD CAPs were used to categorize dHAPs and dLAPs (**Fig. 2**, middle/bottom-right) based on spatial similarity (**Fig. 2**, heatmaps, bottom-left). Both unique and shared dCAPs were identified across high-alpha and low-alpha states. Considering the differences first, for low-alpha, two dLAPs (L1, L2) were spatially similar to visual network BOLD CAPs (B1, B2). This shows that, consistent with prior findings, attenuation of alpha power was accompanied by coactivation of SM, DAN, VAN, and visual networks^10–19^. For high-alpha, two dHAPs (H3, H4) overlapped spatially with the DMN-CON BOLD CAP (B4), suggesting more frequent coactivation of DMN and control networks in the high-alpha state than low-alpha state. The latter had only one dCAP in this group (L4). It is notable that these results are consistent with previously reported patterns of sFC for negative^10–14^ and positive^14–19^ associations respectively. However, these results also show spatially richer patterns consistent with broad and dynamic coactivations associated with alpha power that may vary with temporal context.

Finally, BOLD CAPs (B3, B5), were spatially similar to dHAPs (H3, H5) and dLAPS (L3, L5), indicating activation not unique to low-or high-alpha states. These CAPs encompassed coactivation of SM and attention networks (DAN/VAN) without the visual networks (B3, H3, L3), as well as activation of DMN without control network coactivation (B5, H5, L5). Subtle spatial differences within these patterns may indicate some reconfiguration during high-alpha versus low-alpha states. For instance, considering the DMN BOLD CAP (B5), we see that in the dHAP (H5) instantiation of this CAP there is additional salience network activation, but this is absent in the corresponding dLAP (L5).

### High-alpha state dCAPs differ across datasets

To validate the discovery results, we replicated the entire analysis in an independent publicly available dataset (*n* = 33, Dataset 2^41–43^, **Fig. 3**). Once again, the reference BOLD CAP analysis revealed five dominant CAP states (**Fig 3**. top-right), with functionally similar but non-identical spatial patterns as Dataset 1 **(Fig. 2**; see **S Fig 11** for phase-scrambled results and see **S Fig 12** for results without global signal regression). These could again be classified into visual-network, attention-network, and DMN spatial groups. Two BOLD CAPs were associated with visual network activation: (B1) coactivation of visual and auditory cortices (“visual”, 21%), and (B2) coactivation of visual and the DAN (“visual-DAN”, 18%). Two BOLD CAPs captured attention networks without visual coactivation: one with only the SN (B3, “SM/DAN/VAN”, 19%), and one that included SN and DAN coactivation (B4, “DAN/VAN”, 21%). Finally, one BOLD CAP captured DMN plus control network coactivation (B5, “DMN-CON”, 21%). Thus, these networks were spatially analogous to those identified for Dataset 1.

**Figure 3:**
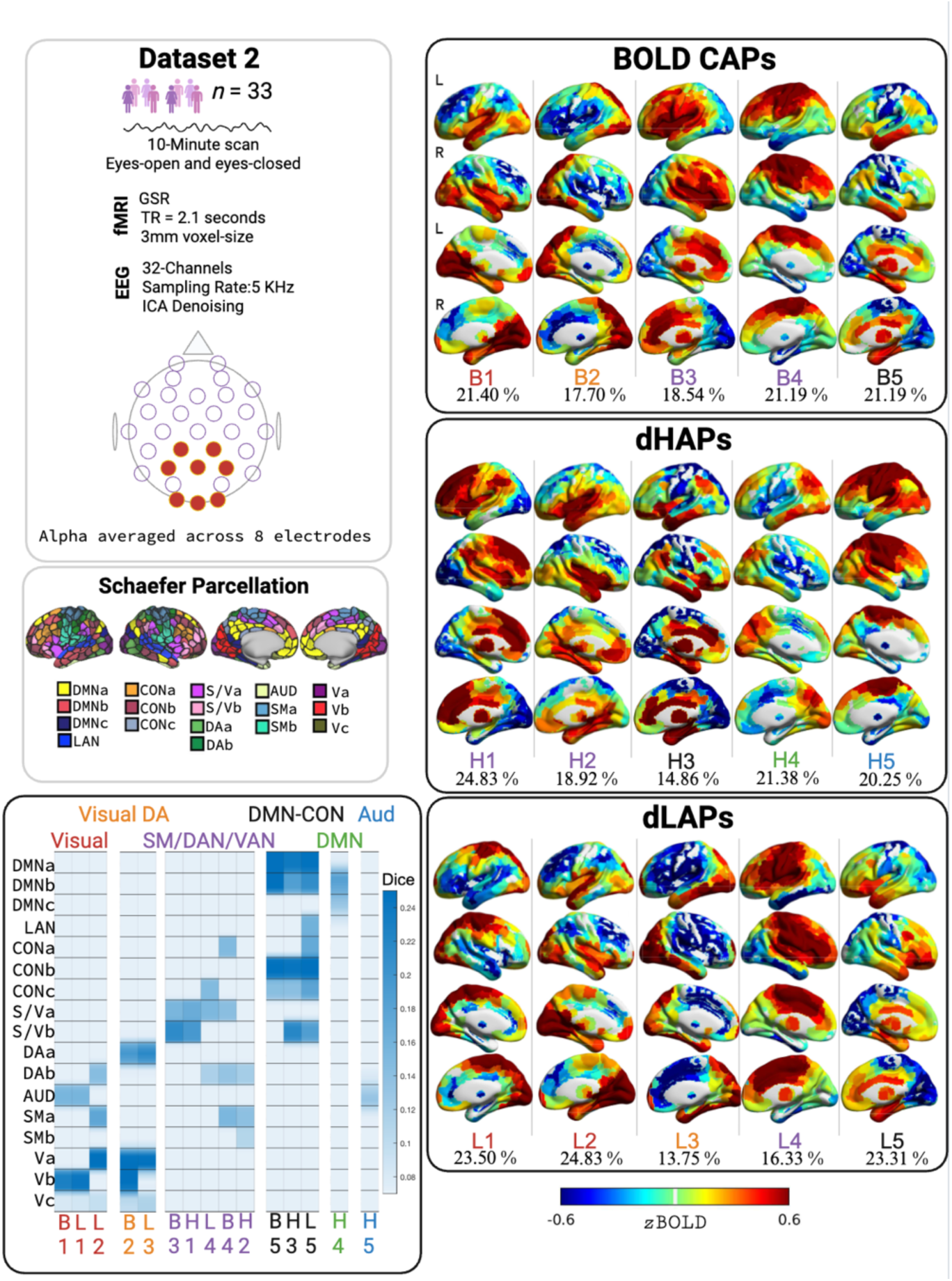
Dataset 2 results. Top Left: Acquisition, preprocessing, and demographic data for Dataset 2. **Middle Left**: Schaefer 400 ROI Kong 17 Network Parcellation^35^ was used to identify canonical BOLD network activation using spin permutation tests and Dice Coefficients. **Bottom Left**: Dice Coefficients from spin permutation tests indicating the overlap between BOLD CAPs and network assignment from the Schaefer Parcellation. Coefficients are only shown when surpassing a threshold of *p* < 0.05. **Top Right**: BOLD CAPs not associated with alpha power identified by using k-means clustering on all BOLD TRs. The frequency that each CAP occurs is shown in percentages. **Middle Right:** Dynamic high-alpha coactivation patterns (dHAPs). **Bottom Right:** Dynamic low-alpha power coactivation patterns (dLAPs). GSR = global signal regression; DMN = default mode network; LAN = language network; CON = control network; S/Va = salience/ventral attention networks; DA = dorsal attention network; AUD = auditory network; SM = somatomotor network; V = visual network; a/b/c = subnetwork a, subnetwork b, subnetwork c; B1, B2, etc. = BOLD CAP1, BOLD CAP2, etc.; H1, H2, etc. = dHAP1, dHAP2, etc.; L1, L2, etc. = dLAP1, dLAP2, etc.

Again, we used these BOLD CAPs to categorize Dataset 2 dHAPs and dLAPs in terms of uniqueness (thresholded at 15% and shown in **Fig 3**. middle-right and bottom-right; (for 10% threshold see **S Figures 13-14**; for 20% threshold see **S Figures 15-16**; for phase scrambled results see **S Figures 17-18**; for results with no global signal regression see **S Figures 19-20**)). Like in Dataset 1, only dLAPs (L1, L2, L3) were associated with visual network BOLD CAPs (B1, B2). These three dLAPs included visual-auditory (L1), visual-SM (L2) and visual-DAN (L2, L3) coactivations. Thus again, decreases in alpha power were associated with visual network coactivations. The dLAPs in this dataset were uniquely associated with DMN (H4), auditory (H5) and DAN/SM (H2) coactivations. Thus, across datasets, we found a commonality of coupling between alpha increases and sub-regions of the default mode network, but also variability in the spatial make-up of these associations. As with Dataset 1, Dataset 2 showed no differences in the frequency of occurrence between dHAPs, dLAPs, and BOLD CAPs (χ²(8, *n* = 299) = 3.53, *p* = 0.90, Cramér’s V = 0.08).

Finally, as for Dataset 1, some dHAPs and dLAPs in Dataset 2 showed similar spatial topography, as referenced to BOLD CAPs. These included SN coactivations (B3, H1, L4) and DMN-control network coactivations (B5, H3, L5), though again with subtle spatial differences. In sum, across datasets, we found that alpha decreases were reliably associated with dCAPs that included visual-coactivation patterns, whereas high-alpha states varied in associated dCAPs that included DMN-control network patterns for Dataset 1, but DMN and auditory networks for Dataset 2.

Low-alpha state CAPs overlap with and expand on static functional connectivity.

The results of the static FC analyses were consistent with previously reported alpha-activations. Unthresholded spatial maps show that alpha power was associated negatively with BOLD signals in the visual cortex, and positively with BOLD signals across intraparietal sulcus (extending into inferior parietal lobule), superior frontal gyrus, and cingulate gyrus for both Dataset 1 and Dataset 2 (**Fig 4**: Top Panel). Dataset 2 shows slightly greater SM negativity compared to Dataset 1 (for results comparing with and without global signal regression see **S Figures 21-24**). Cluster-corrected results (**Fig 4**: Top Panel; *p* < 0.05 voxel-wise uncorrected, *p* < 0.05 cluster-wise corrected) show positive and negative associations remain significant in both datasets.

**Figure 4:**
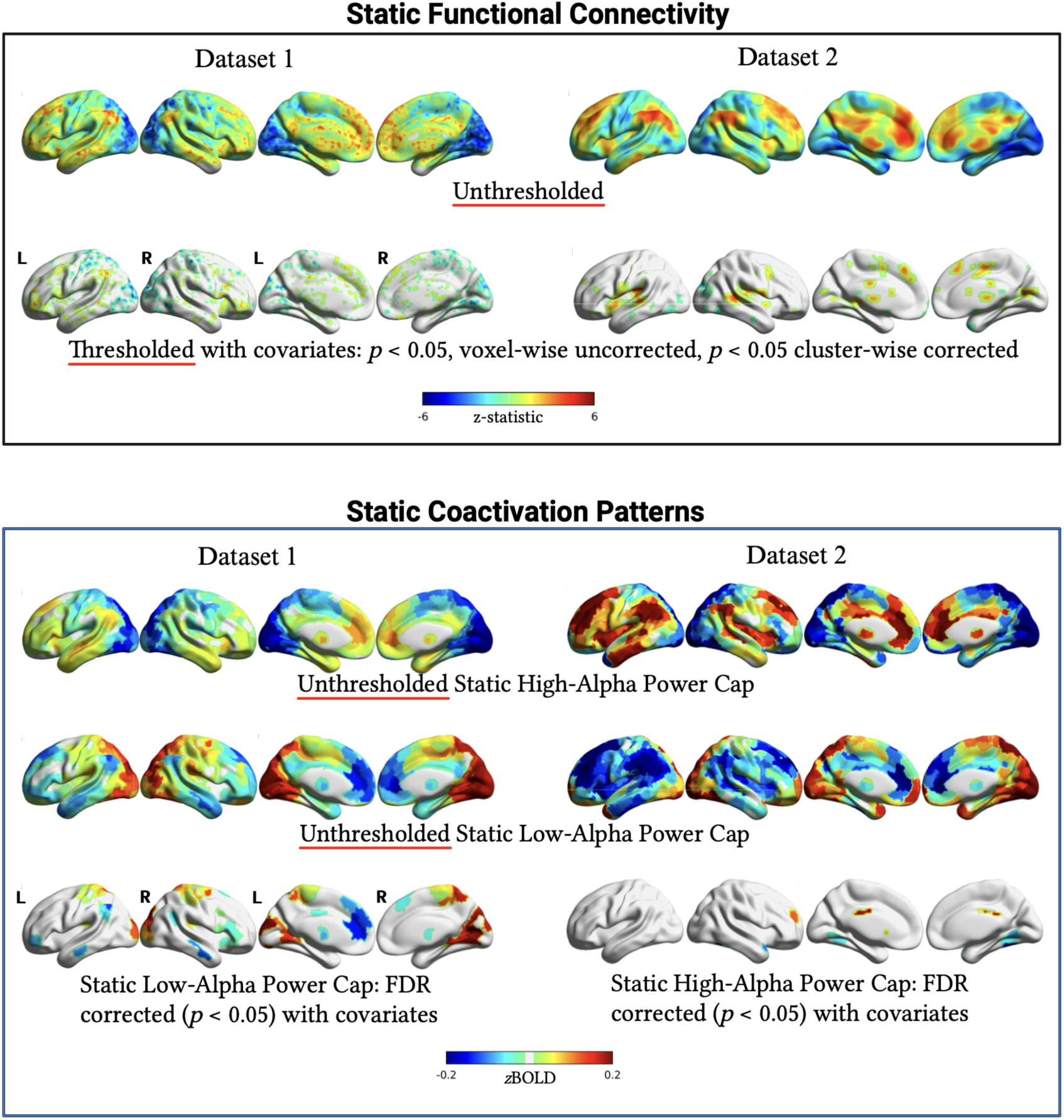
Static functional connectivity and static coactivation pattern analysis. Static functional connectivity. Unthresholded voxel-wise connectivity maps between alpha and the BOLD signal for Datasets 1 and 2 show that alpha is negatively related to BOLD signals in the visual cortex but positively related to BOLD signals within the rest of the cortex. These relationships remain after cluster-correction in a regression model accounting for demographic and scan covariates (Dataset 1: head motion, sex, age, ADHD diagnosis (yes/no); Dataset 2: head motion, sex, age). **Static Coactivation Patterns:** Unthresholded high-alpha and low-alpha power coactivation patterns (CAPs) show similar relationships between alpha and the BOLD signal as the static FC results. During periods of high-alpha power, BOLD activity is suppressed within the visual cortex while other cortices show positive activation. This relationship is reversed during low-alpha power where the visual cortex shows BOLD activity while other cortices show suppressed BOLD activity. These relationships remain in visual, temporal, frontal, and somatomotor cortices when accounting for the covariates used in the static FC analysis (*p* < 0.05, FDR corrected). However, they only remain for the low-alpha CAP in Dataset 1 and the high-alpha CAP in Dataset 2.

This result was replicated using a static CAP analysis (**Fig 4**: Bottom Panel) with time series from the Schaefer 400 ROI Kong 17 network parcellation^35^. Static CAPs were calculated by averaging the BOLD signal for either the top or bottom 15% of TRs of alpha EEG power. Unthresholded static CAP results show the same pattern as sFC results (for results without global signal regression see **S Figure 25**). The high-alpha power static CAP shows negative visual activation in addition to positive activation throughout the rest of the cortex. Accordingly, the low-alpha power static CAP shows positive visual activation in addition to negative activation throughout the rest of the cortex. Thresholded results (FDR correction, *p* < 0.05) from Dataset 1 showed the expected negative relationship between alpha EEG power and the BOLD signal for visual areas and positive relationship with other areas of the cortex. Dataset 2 showed a similar pattern only for the top 15% of thresholded TRs. There were no ROIs that survived FDR correction for the top 15% for Dataset 1 and bottom 15% of Dataset 2.

Thus, the sFC and static CAP analyses showed results consistent with one another, and with prior sFC reports, but captured only a subset of dCAP findings. The static results replicate previous reports that alpha generally has a negative relationship with BOLD activity in the visual cortex^44–46^. However, they did not reliably differentiate canonical network coactivation outside of visual cortex as in the dCAP analyses (**Figs. 2, 3**). This implies that low-alpha states may be associated with more distributed coactivations than typically reported. These dLAPs vary in coactivation outside of visual cortex (i.e., involving SM, VAN and DAN) across time series and datasets making them difficult to detect using static analyses that may collapse across this variability to only show coactivation predominantly in visual cortex. Similarly, the positive associations in the static results corresponded to only a subset of the dHAPs observed across datasets. Putatively, given the variability of dHAPs, the static analyses may not be sensitive to their transient activation.

### Alpha-state CAPs are differentially associated with anterior and posterior thalamus

A common finding in alpha-power EEG-fMRI studies is a positive association with thalamic BOLD signal, in opposition to negative association with cortical BOLD signal^11,21,29,32,44^, with a proposed role of thalamus in alpha generation. However, this finding has been somewhat perplexing, as posterior thalamus is often functionally connected with the visual cortex and thus would be expected to follow the same (not opposite) direction of association with alpha power. A potential explanation was proposed by Liu et al., in 2013^47^, who used ultra-high field mapping of thalamus along with concurrent EEG-fMRI to show that the nature of BOLD-alpha correlation varies with location in thalamus – positive (as previously reported) for the anterior and medial thalamus but negative (consistent with visual cortex coactivation) for the posterior thalamus. They proposed that the pulvinar and visual cortex form a common alpha-related network during visual processing, whereas the anterior and medial dorsal nuclei belong to a separate network involved in vigilance and arousal.

As each dCAP included subcortical ROIs, we examined this possibility and qualified subcortical contributions to BOLD CAPs, dHAPs, and dLAPs. First, we considered the dLAPs consistently associated with visual cortex BOLD signals. The corresponding visual network coactivation patterns (L1, L2) included positive signals in posterior thalamus but negative across anterior thalamus (also present in the corresponding BOLD CAPs B1, B2) (**Fig. 5)**. In complement, in Dataset 2 (**Fig. 5**), visual-network BOLD CAP (B2), and dLAP (L2), also included stronger activity in posterior-compared with anterior-thalamus. This is consistent with Liu et al., 2013^47^, whereby posterior thalamus is negatively associated with alpha power as with visual cortices. In complement, the most unique high-alpha dCAP in Dataset 1 (H3), was associated with broad increases in signal across anterior thalamus, along with the DMN. This pattern was also true for dHAP (H3) (DMN-CON) and dHAP (H1) (SN) in Dataset 2, supporting the proposed association of anterior thalamic activity associated with regulatory, or vigilance related networks.

**Figure 5:**
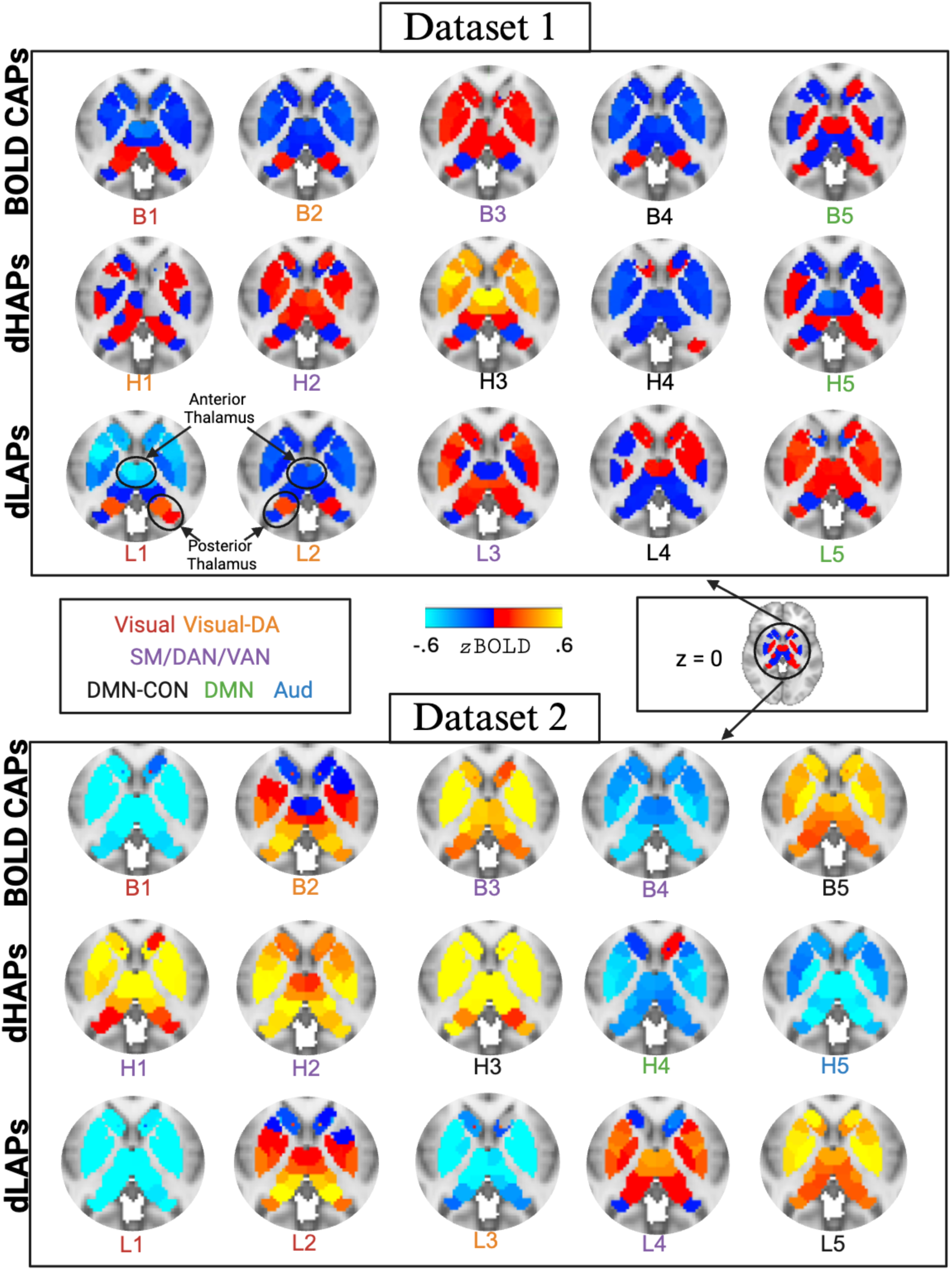
Subcortical activation from BOLD CAPs, dHAPs, and dLAPs, represented using 54 ROIs from the Tian parcellation. Top: Dataset 1 shows stronger thalamic activation in posterior areas compared to anterior areas in BOLD CAPs 1 and 2 and well as dLAPs L1 and L2. These accompany visual cortical activation from Figure 2. **Bottom:** Dataset 2 shows similar patterns stronger activity in posterior compared to anterior thalamic ROIs in BOLD CAP 2 and dLAP L2.

## Discussion

The current study identified occipital EEG alpha oscillations associated with dynamic BOLD brain states. The use of a dCAP analysis with two concurrent EEG-fMRI datasets, facilitated identification of transient coactivation patterns during both low-and high-alpha intervals during resting-state. Several low alpha dCAPs replicate prior sFC results showing increased visual network activity but also present with additional coactivation within sensorimotor, auditory, and attention networks. However, multiple coactivation patterns for both high and low alpha periods overlap across canonical networks (DMN, CON, SN, DAN, VAN, SM) as well as subcortical activations. These results suggest that visual, SM, auditory, and attention network coactivation was generally associated with low-alpha power, while coactivation of networks such as the DMN and CON, involved in internal and external cognition^48^, were generally associated with both low-and high-alpha power. These findings reveal that multiple network configurations produce similar alpha modulation patterns, supporting a temporal-or context-dependent dynamic network framework for the gating function of alpha oscillations.

These multiple configurations are broadly anchored in two functional states corresponding to high-alpha and low-alpha occipital cortical excitability^1–3^. During low-alpha states, excitability is documented to increase in cortex^10–14^. Our results support and expand on this finding. Across datasets, dLAPs were consistently associated with visual network BOLD activation in addition to SM, auditory, and attention network coactivation. Thus, decreases in alpha are consistently associated visual and attention network coactivation that sometimes include sensorimotor and auditory networks. This result is notable, given vast differences between datasets in acquisition and signal processing parameters, thus offering a robust replication. Along with convergent findings in sFC and static CAP analyses of low-alpha states, and consistency with prior report^10–13^, our finding support the reliability of the association and the robustness of the proposed analytic approach. Expanding on prior reports, the visual dLAPs also showed heterogeneity across datasets, including the SM and DAN in Dataset 1 but more localized to posterior cortex in Dataset 2. These results echo prior reports that alpha oscillation reduction over posterior occipital scalp track visual system BOLD activation but also extend into associative cortices in some contexts^10–14,29^. This implies that decreases in occipital alpha power reflect multiple network interactions of visual cortices rather than a static, singular network – an idea that resonates with proposals for multiple cortical alpha sources^49,50^.

This heterogeneity becomes emblematic across both low-and high-alpha states when focusing on coactivation of DMN and CON networks. Several dLAPs and dHAPs involve the previously reported CON^10,11^, DMN^14–17^, and SN (also called cingulo-opercular network)^18,19^, and included novel observations of auditory cortex engagement. While the CON network typically shows robust task-based activity in fMRI studies of executive function^51^, the DMN still plays a role in updating task-context involving task rules or experimental organization^52^. Further, communication between the DMN and CON network plays an important role in social processing that also requires the ability to incorporate internal (e.g., awareness of emotional states) and external information (e.g., awareness of the emotional states of others)^53^. Thus, the interplay between the DMN and CON networks are important for integrating internal and external information for social and goal-driven behavior. The interesting finding is that the activity of the DMN and CON networks generally did not depend on alpha power. This contrasts with the results suggesting that visual network BOLD activation is mainly found during periods of low-alpha power. Previous research showing that BOLD activity can be uniquely associated with alpha, beta, and gamma oscillations in resting-state and task-fMRI designs^54,55^ could explain why CON and DMN coactivation is not always associated with low-or high-alpha power. The current study focused on occipital alpha oscillations during resting-state, and it will be important to identify how different sources of alpha, beta, and theta in various task-fMRI contexts may be associated with BOLD CON and DMN coactivations.

These results may help reconcile findings from previous resting-state EEG-fMRI studies by demonstrating that multiple states can co-exist within the same scan session, with different states dominating a given recording depending on the temporal context. In such cases, sFC analyses would only identify networks contributing most strongly to the covariance structure across all time points. As such, prior interpretations of increased alpha power as indexing internally directed processes^14–17^ or a state of tonic alertness (without internal mentation)^18,19^ can both hold true. In complement, increased alpha power has been linked to reduced cortical excitability, across a variety of contexts that include suppression of distracting visual inputs^7^, as well as passive states when cortical demand is low – such as post task-cessation^56^ or the so-called “idling” state^57^. Collectively, the data suggest that alpha oscillations are a domain-general gating mechanism on gating cortical excitability (in the present case converging on visual cortex), triggered through multiple neural configurations^26^.

Several important caveats merit mention regarding dHAP interpretation. First, the association between high-alpha intervals and non-visual networks was not exclusive to dHAPs, though nuanced differences existed between dHAP and dLAP instantiations. For instance, the DMN network in H5 vs L5 included the SN in the former but not the latter, potentially indicating meaningful differences about the functional state of the network related to alpha oscillations. Second, the current results suggest that visual cortex activity is the central target of cortical gating via reduced activation in the presence of alpha oscillation. This result may be related to analysis of occipital-electrode alpha power, where visual source contributions are likely to be strongest. Alpha range (8-12 Hz) oscillations have been documented in motor^57^ and auditory^58^ cortices and may show similar effects. Finally, the present findings are independent of the proposed role of alpha oscillations in supporting long-range synchronization among cortical regions^18,59,60^, though are not in contradiction with those frameworks.

An important and unexpected finding was the thalamic contribution to alpha-state CAPs. We identified a dissociation between posterior and anterior thalamic nuclei, with posterior thalamus showing negative associations with alpha power together with the visual cortex, and anterior thalamus showing associations with both low-and high-alpha power together with the DMN, SN and CON networks. This pattern is consistent with Liu et al., in 2013^47^, who reported opposing alpha-BOLD relationships across thalamic subdivisions based on concurrent EEG-fMRI and high field anatomical and FC mapping thalamus. Additionally, the dissociation was observed more robustly in the eyes-open Dataset 1, and less reliably in Dataset 2 in which participants could choose to close or open their eyes during the recording. This falls in line with posterior thalamocortical circuits being preferentially engaged during visual processing, and with significant reconfiguration in alpha-BOLD coupling with elimination of visual input^29^. In complement, the anterior and midline thalamic nuclei may contribute to the regulation of large-scale cortical state through their positive coupling with alpha oscillations and transmodal networks^47^. Broadly, this positive association aligns with reported coupling of alpha power with physiological signals^22^, such as respiration^23^, though whether these associations reflect arousal-based modulation of thalamo-cortical activity, a thalamo-oscillatory correlate of canonical network anti-correlations^48^, or is an independent contribution to the alpha signal, remains to be determined.

Finally, the present results emphasize the value of a dynamic approach to analysis of oscillatory network correlates^28–31^, while also offering insight on potential strengths versus weaknesses of dynamic versus static analyses. In particular, the sFC analysis yielded fewer statistically significant effects for Dataset 1 which had a shorter recording duration (6 minutes) than Dataset 2 (10 minutes), despite the former including more participants. This observation is in line with prior recommendations for resting-state FC analyses^61^. In complement, the reliability of the conclusions across datasets suggests that dynamic analysis may be less susceptible to this limitation, leveraging total data length across sample size and scan duration. The subcortical results, in complement, show sensitivity to sub-thalamic differences potentially obscured in sFC analyses of alpha power. Finally, it is perhaps notable that none of the results – BOLD CAP, dLAP nor dHAP – showed states characterized by a single canonical network but rather combinations of these networks, consistent with a dynamic process with brain networks reconfiguring transiently on a timescale of seconds^37,62^

In summary, this study provides evidence across two independent datasets that EEG occipital alpha oscillations are associated with dynamic BOLD brain states. Low-alpha states are robustly associated with state-specific BOLD activation of visual cortex, SM, attention networks in addition to the posterior thalamus. In contrast, though high-alpha states are associated with decreased cortical BOLD activation across visual networks, this occurs alongside state-independent (they also occur for low-alpha) DMN, CON, and attention network coactivation in addition to the anterior thalamus. These results suggest a dynamic framework of alpha-BOLD associations, that address unresolved heterogeneity across previous sFC EEG-fMRI studies and highlight the breadth of the gating function of alpha oscillation in network dynamics.

## Acknowledgements

This work used computational and storage services associated with the Hoffman2 Cluster which is operated by the UCLA Office of Advanced Research Computing’s Research Technology Group. We also thank Timothy Kelley for help with data acquisition.

## Data Sharing

The raw data for Dataset 1 is available at: https://nda.nih.gov/edit_collection.html?id=3101

## Methods

### Participants

#### Dataset 1

A total of 94 subjects were included in the CAP analysis while two smaller subsets of participants with available behavioral assessments of ADHD symptom severity for the KSADS and SNAP inattention and hyperactivity subscales along with a total score measure (**S Table 1**). This sample was part of a larger longitudinal study investigating attentional control and spatial working memory in individuals with and without ADHD (n = 176). Eligibility was based on participation in prior research within the center and between 14 and 39 years of age, with exclusions including: history of neurological illness or injury; diagnosis of suicidality, psychosis, mania, significant aggression or substance abuse; inability to discontinue ADHD medication (24 to 48 hours prior to session based on half-life of medication); inability to undergo neuroimaging exams (either EEG or MRI) including due to metal in body, pregnancy and claustrophobia. Individuals under the age of 18 provided assent in addition to a parent or guardian’s consent. Participants received $150 as compensation for their time. All methods were performed within the relevant guidelines and regulations in accordance with the Declaration of Helsinki, and in accordance with the approved protocols by the Human Research Protection Program of the University of California, Los Angeles. There were 105 subjects with both fMRI and EEG resting-state data. Due to data loss during preprocessing and quality control (e.g., poor registration) and data quality criteria for the CAP analysis (subjects with alpha EEG time series shorter than 50% of the rest-scan were removed), the final sample size was 94 subjects (Mean age = 23.85 years old, SD = 5.17; range = 17-39 years of age; 37 females).

#### Dataset 2

The second dataset consisted of publicly available multi-modal EEG-fMRI data^41–43^ for 33 subjects (Mean age = 22.1, SD = 3.2; range = 18-30 years of age; 17 females) consisting of a single 10-minute resting-state scan, whereby participants could choose to keep their eyes closed or open.

### Clinical & Neurocognitive Assessment

#### Dataset 1

Participants completed both a clinical interview performed by a clinical psychologist, as well as a neurocognitive assessment. The full set of measures collected is provided in supplemental materials and will be the focus of future publications. Relevant to the current project are inattention symptoms, quantified via total number of symptoms met during diagnosis (clinician assessed). The symptoms were used here as a metric of individual differences to assess behavioral impact of neural features on individual differences in attention.

### Neuroimaging Session Protocol

#### Dataset 1

Following consent & clinical assessments, participants completed a single neuroimaging session that lasted 2-3 hours. The session included an introduction and safety screening for the MRI portion of the session. This was followed by an EEG-only practice session (30 min), and a concurrent EEG and fMRI session (90 min). The eyes-open rest data used in the current analyses were acquired as part of the latter, along with several cognitive tasks including a Sternberg spatial working memory task (SWM)^63,64^, one run of alternating eyes-open eyes-closed rest (EOEC) and one run of a passive viewing task (PV). As they are not the focus of the present analyses, the tasks are fully described in supplemental materials. However, performance on the spatial working memory task, which include accuracy and reaction time variability, were used here as a behavioral correlate of individual differences related to identified CAPs.

### fMRI Image acquisition

#### Dataset 1

MRI data were acquired on a 3T Siemens PRISMA MRI at the UCLA Staglin Center for Cognitive Neuroscience using a 20-channel (functional scans) and a 32-channel (structural) head coil. Following localizer and field mapping, we acquired five functional scans, which used a Siemens product echo planar imaging sequence (TR = 1000 ms; TE = 25 ms; flip angle = 70°; FoV = 192 mm; matrix = 64×64; number of slices = 56; slice thickness = 2.5 mm (0.5 mm gap), slice acceleration factor PE = 1). The complete acquisition included: two runs of a SWM task, followed by eyes-open/eyes-closed and passive viewing control conditions (see Supplemental Materials for details). The 6 min eyes-open rest scan used in the current CAP analyses was acquired next. Following the functional scans, we switched to a 32-channel structural coil and removed the EEG cap to ensure that participants are comfortable and still for the subsequent anatomical scans. Aw high-resolution T1-weighted MPRAGE image was used for registration (TR = 1900 ms; TE = 2.48 ms; flip angle = 9; FoV = 256 mm; slice thickness = 1 mm; slices = 208).

#### Dataset 2

MR data were acquired on a 3 Tesla Siemens Prisma Fit scanner using a 20-channel receive-array coil at Pennsylvania State University. Structural MRI data were acquired using a magnetization-prepared rapid acquisition gradient echo (MPRAGE) sequence with the following parameters: TR = 2300 ms, TE = 2.28 ms, TI = 900ms, the axis of acceleration: phase encoding direction along the A-P axis of the brain, flip angle = 8 degrees, FOV = 256 mm, matrix size = 256 x 256 x 192, voxel size = 1mm x 1mm x 1mm, acceleration factor = 2. Blood oxygen level dependent (BOLD) functional MRI data were acquired using an echo planar imaging (EPI) sequence with the following parameters: TR = 2100 ms, TE = 25 ms, flip angle = 90 degrees, slice thickness = 4 mm, number of slices = 35, FOV = 240 mm, in-plane resolution = 3mm x 3mm. The rest scan was 10 min in duration and participants could choose if they wanted to keep their eyes open or closed.

### fMRI Preprocessing

#### Dataset 1

FMRIB’s Software Library (FSL, v.5.0.1)^65^ was used for preprocessing of functional images. First, the brain was isolated from the surrounding tissue using BET. Then, to correct for subject motion, functional images in each block were realigned to the middle volume by applying a rigid body (6 degrees of freedom) transformation using a normalized correlation similarity function with trilinear interpolation. The motion parameters from each transformation were retained for subsequent analysis. Data were smoothed spatially using a 5 mm full-width-half-maximum Gaussian kernel, filtered temporally using a non-linear high-pass filter with a 90 s cut-off. The data were then denoised by means of probabilistic independent components analysis^66^, with automatic classification of artifact components using a hierarchical supervised classification framework (ICA-FIX)^67^ custom trained on in-house data (accuracy in classifying noise components at 90.47%). Identified artifact ICs were subtracted from the data. To establish the efficacy of this two-step motion removal approach we re-ran the motion correction realignment algorithm on the ICA denoised data and evaluated change in motion and framewise statistics. Specifically, we extracted the derivative of root mean square of variance (DVARS) and framewise displacement (FD) values^68^ for each run and for each subject, before and after denoising the data with ICA. Across all runs, effects of diagnostic group, age, and gender and their interaction with time (pre/post cleaning) were not significant, whereas the effect of cleaning (pre/post) was significant (*p* <.02, FD_pre_=.062mm, FD_post_=0.036mm, DVARS_pre_=55, DVARS_post_=43). The functional images were then registered to the high resolution T1 scan and then to the standard MNI (Montreal Neurological Institute) space (2mm) using affine transformations from FSL. Finally, the global signal (GS) was calculated using a whole-brain SPM mask and regressed out of the data. GS regression after ICA-FIX has shown to mitigate the impact of head motion on the BOLD signal compared to either process alone^69^.

#### Dataset 2

These data were originally used in Phadikar et al. (2025)^28^, and thus full processing steps are described therein. Preprocessing was done using SPM12 and AFNI tools. The 10-minute resting-state scan was first despiked using AFNI then underwent realignment, spatial normalization to MNI space (3mm), and smoothing (FWHM at 6mm). The GS was calculated using a whole-brain SPM mask and then regressed out of the data.

### EEG acquisition

#### Dataset 1

The EEG data were acquired using Electrical Geodesics (Eugene, OR) NS300 (EEG-only, running NetStation 5.4.1.2 software) and NS400 (EEG-fMRI, running NetStation 5.4.2 software) system, the latter sampling at 1000Hz and clock synchronized to the MR EPI acquisition. We used 128-channel high-impedance HydroCel Geodesic Sensor Nets, with reference at vertex. Nets were positioned on each participant by aligning the vertex electrode with the vertex of the head, identified at the midpoint of the anion-to-inion and left/right preauricular landmarks. Electrode impedances were <50 kΩ, as recommended by the manufacturer. Two additional external electrocardiogram (ECG) electrodes were applied to the chest (positions: left midclavicular line in 5^th^ intercostal space and 4^th^ left intercostal space at sternal border) for later use in suppression of artifacts as described below.

#### Dataset 2

For this publicly available dataset, the EEG acquisition parameters are in published references^41–43^.

### EEG Preprocessing

#### Dataset 1

Offline EEG processing and analyses were performed using custom MATLAB (R2023, The MathWorks, Inc.) scripts using functions from the EEGLAB environment (v.2023)^70^. The EEG data were high pass filtered (>.1 Hz). Gradient-related artifacts generated during each TR of scanning were removed next, using in-house scripts that first, for each EEG sample within a TR, modeled and removed linear drift to remove amplitude signal and then used a moving window (13 TRs centered) average template subtraction^71^ to eliminate gradient artifact. The time series were then downsampled to 250Hz and trimmed to remove residual gradient artifacts at the beginning and end of the time series. We additionally removed ballistocardiogram effects^72^ in two steps. First, we used a template-based approach^73^ to identify and then subtract the mean BCG artifact from each BCG event. However, here we used an adaptation of this approach^74^ that seeks to identify the heartbeat events in the EEG data (i.e., BCG events) rather than in the EKG data (i.e., EKG events) as typically reported^63^. This is because we have previously found the timing between EKG and BCG peak events to be variable across individuals and not consistent with published recommendations^72^. In contrast, the timing of the BCG relative to the ECG events within an individual is highly reliable and thus provides at least as good if not better estimate of the BCG artifact^75^. In the second step the BCG-template cleaned data were decomposed into maximally independent component (IC) processes by temporal ICA decomposition using extended infomax^76^ (stopping weight = 1e-7, maximum learning steps = 1000). We identified ICs that captured residual BCG artifacts by creating an event-related average for each IC, time-locked to BCG peaks, and then tested for significant root-mean-square signal in the event-related average. Such ICs were subtracted from the data. We used ICLabel^77^ machine learning functions in EEGLAB to identify artifacts due to eye movement and residual high-frequency noise. These ICs were also subtracted from the data. Finally, electrodes with poor signal were identified based on weak correlations with other channels (<.1) for more than 40% percent of the session (flt_clean_channels.m). The identified electrodes were interpolated, and the data were re-referenced to average reference.

#### Dataset 2

Resting-state EEG data were preprocessed using the EEGLAB toolbox in MATLAB. Scanner-related gradient artifacts were removed using the FASTR algorithm implemented in the FMRIB plugin. The data were then downsampled to 1 kHz and bandpass filtered between 1 and 30 Hz. Cardiac artifacts were identified and removed using the FMRIB toolbox, after which the data were re-referenced to the common average. Finally, independent component analysis (ICA) was applied to decompose the EEG into independent components, from which residual pulse, muscle, and eye movement artifacts were identified and removed.

### EEG Feature Extraction

#### Dataset 1

To quantify modulation of alpha power, the continuous EEG time series were segmented into non-overlapping contiguous 2 sec epochs. This epoching approach was selected to facilitate artifact removal procedures. We additionally removed any epochs containing residual artifacts, detected automatically using statistical thresholding (pop_autorej.m, startprob = 5, maxrej = 3%). This procedure produced a maximum of 180 possible epochs per subject. To calculate event-related alpha power modulation, we applied the fast Fourier transform function, as implemented in Matlab, to the epoch time series. For each epoch, total power in the alpha frequency band (8-12Hz) was extracted using the bandpower function, which integrates power within a frequency range of a power density function using the rectangle method. These values were divided by the total power (across all frequencies in 1-40Hz range) in that epoch. This epoch normalization was included to account for large changes in total power (due to changes, for example, in residual noise in the EEG time series from MR or BCG artifacts) across epochs. Finally, the normalized alpha power values were additionally averaged across electrodes to reduce the dimensionality to a single number. These electrodes were selected based on the distribution of the difference in alpha modulation during eyes-open versus eyes-closed blocks, which produce a known large-effect modulation of occipital alpha oscillation. We identified the top 20 electrodes showing a significant difference (E52, E59, E60, E65, E66, E67, E70, E71, E72, E74, E75, E76, E77, E83, E84, E85, E89, E90, E91, E92), and these correspond to posterior scalp surrounding electrodes O1, Oz, O2 in the 10-20 system.

For use in CAP analyses, several additional steps were implemented. As each participant’s average alpha power time series was calculated at a rate of 2 seconds per measure, in order to match the acquisition rate of the BOLD signal that was acquired at 1 second per TR, the alpha time series was resampled using linear interpolation to rate of 1 second per measure. Due to artifact removal involving censoring, not every participant had an alpha time series that was 360 TRs in length (the length of the BOLD fMRI scan). Any missing time-points in the alpha time series from artifact elimination were also replaced with linear interpolation at this point. Next, the alpha time series was convolved with a BOLD HRF signal modeled from a TR of 1 second from the SPM toolbox. This was done to account for the hemodynamic lag of the BOLD signal relative to the EEG signal. Additionally, the first 15 TRs from the alpha-EEG signal were eliminated to account for the effect of the HRF convolution at the start of the time series. The first 15 TRs from the BOLD data were also eliminated to match the alpha-EEG time series length. This resulted in a maximum possible length of 345 TRs for analysis.

#### Dataset 2

Relative alpha power (divided by total power in extracted frequency range 1-40 Hz) was calculated using non-overlapping 2100 ms windows (i.e., one BOLD TR) with a Hamming taper. For each window, we removed DC (detrending), computed the single-sided power spectral density (PSD), and then integrated the PSD over 8–12 Hz (sum × df). Alpha power was then averaged across eight posterior electrodes: O1, O2, Oz, P3, P4, CP1, CP2.

### Static Functional Connectivity Analysis

#### Datasets 1 and 2

To replicate previous research showing an inverse relationship between BOLD signal intensity and the alpha-EEG time series within the visual cortex, a static functional connectivity analysis was conducted. The alpha-EEG time series from each subject was associated with each voxel in the brain using linear regression. One-sample t-tests were then conducted on the voxel-wise whole-brain beta estimates (*p* < 0.05 voxel-wise uncorrected and *p* < 0.05 cluster-wise corrected) with and without nuisance regressors. Dataset 1 regressors included age, sex, head motion (framewise displacement), and ADHD diagnosis (yes/no). Dataset 2 regressors included age, sex, and head motion.

### Coactivation Pattern Analysis (CAP)

#### Datasets 1 and 2

Two types of CAPs were identified: static and dynamic. Static CAPs were identified to validate that the increased level of BOLD signal activation from previous research in the visual cortex was associated with the lowest points of alpha EEG intensity. Static CAPs were calculated using the Schaefer 400 ROI 17 network parcellation in order to reduce computational demands. The average BOLD signal for each ROI was associated with relative alpha EEG power for the top or bottom 15% TRs of activation were then averaged together. The 15% threshold is based on previous research showing that this level of thresholding produces spatial maps that are correlated with sFC spatial maps at *r* >.99^33^. This produced two sets of CAPs: one representing average BOLD activation when alpha power is highest and one representing average BOLD activation when alpha power is lowest. One sample *t*-tests were conducted for each ROI using the same nuisance regressors from the sFC analysis (*p* < 0.05, FDR corrected)^78^.

Dynamic CAPs were used to identify different configurations of BOLD activation generally associated with the strongest and weakest TRs associated with alpha EEG intensity. Dynamic CAPs were also calculated using the Schaefer 400 ROI 17 network parcellation. The average time series across all voxels within each ROI was calculated for all subjects. Next, the top and bottom 15% of TRs representing the highest and lowest intensity of the alpha EEG signal was identified for each subject. The top and bottom TRs represent the “peaks” and the “troughs” of the alpha-EEG time series respectively. The BOLD “top” thresholded TRs and the “bottom” 15% of thresholded TRs were then concatenated into a matrix of the top 15% of BOLD TRs related to alpha EEG activation and a matrix of the bottom 15% of BOLD TRS related to alpha EEG activation across subjects. Each matrix was subject to k-means clustering.

The optimal k-means clustering cluster solution was identified using the elbow criterion (**S Figure 26**) by maximizing the ratio consisting of contrasting within-and between-cluster distances across clustering solutions from 2 - 20^37^. Dynamic CAP states were then identified using the optimal k-means clustering (5000 iterations and 5000 replications) solution for both the top and bottom 15% of BOLD TRs related alpha EEG intensity. Finally, the Network Correspondence Toolbox (NCT)^40^ was used to identify the canonical networks associated with each brain state via dice coefficients computed with spin test permutations.

Note that, following this procedure, for Dataset 1 the total number of TRs (duration = 1 sec) included across participants varied due to the availability of usable EEG data. In cases where EEG data was not acquired for the length of the entire run, the fMRI resting-state data was trimmed to match the length of the alpha time series. The average number of TRs included in the 15% threshold CAP analysis was 46.23 (SD = 4.05, range: 27-52) of a possible 52 TRs meaning that the average length of the available alpha-EEG time series was 308 TRs (SD = 27; range: 180-345; subjects with less than 180 TRs were removed from the analysis). All BOLD data was trimmed to match the length of the alpha-EEG time series. Due to different methods in EEG preprocessing, Dataset 2 had no missing EEG frames, and each subject had all TRs for both alpha EEG power and BOLD activation.

### BOLD Coactivation Pattern Analysis and Null Modeling

#### Datasets 1 and 2

In order to ensure that the dynamic CAP analysis related to the average alpha-EEG signal were capturing dynamic network CAPs distinct from overall average BOLD CAPs, k-means clustering was run on all subjects using all TRs using the optimal cluster solution from the dynamic CAP analysis. This was used to test if and which of the network activation patterns associated with the alpha EEG signal were distinct from the dominant patterns of activity in the overall data.

Additionally, null model comparisons were created via phase scrambling of the ROI BOLD time series while preserving the mean, standard deviation, and autocorrelation of the original time series^32,33^. Phase scrambled time series were used in static and dynamic CAP analyses as well as the group k-means clustering. This was used to show that the BOLD activation related to alpha EEG power was unlikely to be composed of random noise and likely represents neural activity.

### Brain and Behavior Associations

#### Dataset 1

To test if measures of ADHD symptom severity and working memory task performance were associated with the frequency of CAP occurrence, regression models were run using the frequency of CAP occurrence as the dependent variable (DV) and measures of ADHD symptom severity as the independent variables (IV) along with nuisance regressors consisting of age, gender, and head motion. ADHD symptom severity was assessed using the SNAP and KSADS questionnaires with the inattention, hyperactivity, and total scores for each measure (**S Table 2**). There were no significant associations between CAP frequency and behavioral scores (at ceiling though), nor symptoms.

