## Supplemental Materials for "Alpha-range neural oscillations are associated with multiple dynamic brain networks"

**Supplemental Table 1**: Demographic information for the overall sample and the two samples used for the brain-behavior associations. i = inattentive; h = hyperactive; c = combined; t = total score; TD=typical development.

**Overall Sample**

| *Sample* | *Mean Age(SD), Range* | *Sex* |
|---|---|---|
| N = 94 | 23.84(5.17) 15.5-40 | 37F |
| TD=41 | 23.90(5.37) 15.5-40 | 17F |
| ADHD-C = 25 | 24.66(4.85) 16.42-32.92 | 9F |
| ADHD-I = 26 | 23.42(5.36) 17.08-39.08 | 11F |
| ADHD-H = 2 | 18.91(0.47) 18.58-19.25 | 0F |

**<u>Supplemental Table 2:</u>** Demographic information for subjects used in brain-behavior analyses from Dataset 1.

**KSADS Sample**

| *Sample* | *Mean Age(SD)* | *Sex* | *KSADS (i)* *Range: 0-9* | *KSADS (h)* *Range: 0-9* | *KSADS (t)* *Range:0-18* |
|---|---|---|---|---|---|
| N = 93 | 23.85(5.17) 15.5-40 | 36F | 4.56(3.67) | 2.77(2.79) | 7.33(6.05) |
| | | | 0-4 | 0-4 | 0-7 |
| 41 TD | 23.90(5.37) 15.5-40 | 17F | .88(1.35) | .41(.92) | 1.29(2.08) |
| | | | 6-9 | 5-9 | 11-18 |
| 25 ADHD-C | 24.58(4.76) 16.42-32.92 | 5F | 8.32(1.07) | 6.48(1.45) | 14.80(1.89) |
| | | | 5-9 | 0-4 | 6-13 |
| 25 ADHD-I | 23.42(5.36) 17.08-39.08 | 10F | 7.04(1.57) | 2.64(1.11) | 9.68(1.77) |
| | | | 1-3 | 6-7 | 7-10 |
| 2 ADHD-H | 18.91(0.47) 18.58-19.25 | 0F | 2(1.41) | 6.5(0.71) | 8.5(2.12) |

**SNAP Sample**

| *Sample* | *Mean Age(SD)* | *Sex* | *SNAPS (i)* *Range: 0-27* | *SNAPS (h)* *Range: 0-24* | *SNAPS (t)* *Range:0-51* |
|---|---|---|---|---|---|
| N = 73 | 25.15(4.96) 15.5-40 | 25F | 11.68(8.06) | 8.23(6.51) | 19.92(13.69) |
| | | | 0-20 | 0-22 | 0-36 |
| 34 TD | 25.02(5.20) 15.5-40 | 13F | 6.20(5.57) | 3.97(4.32) | 10.12(8.81) |
| | | | 3-27 | 5-24 | 10-51 |
| 20 ADHD-C | 25.98(4.16) 18.5-32.92 | 7F | 17.35(7.49) | 13.35(5.26) | 30.70(11.77) |
| | | | 0-27 | 1-19 | 2-45 |
| 19 ADHD-I | 24.51(5.44) 18.33-39.08 | 5F | 15(7.15) | 11.25(6.17) | 26.25(11.85) |

**Task Performance Sample**

| *Sample* | *Mean Age(SD)* | *Sex* |
|---|---|---|
| N = 89 | 23.87(5.25) 15.5-40 | 35F |
| 41 TD | 23.90(5.37) 15.5-40 | 17F |
| 24 ADHD-C | 24.57(4.96) 16.41-32.92 | 9F |
| 23 ADHD-I | 23.31(5.57) 17.08-39.08 | 9F |
| 1 ADHD-H | 18.58 | 0F |

**Supplemental Figure 1**: Dataset 1 dCAPs for high-alpha periods (dHAPs) using a 10% threshold for alpha power. The order of states is arbitrary and does not necessarily match the main manuscript ordering. The frequency that the state occurs is shown in percent.

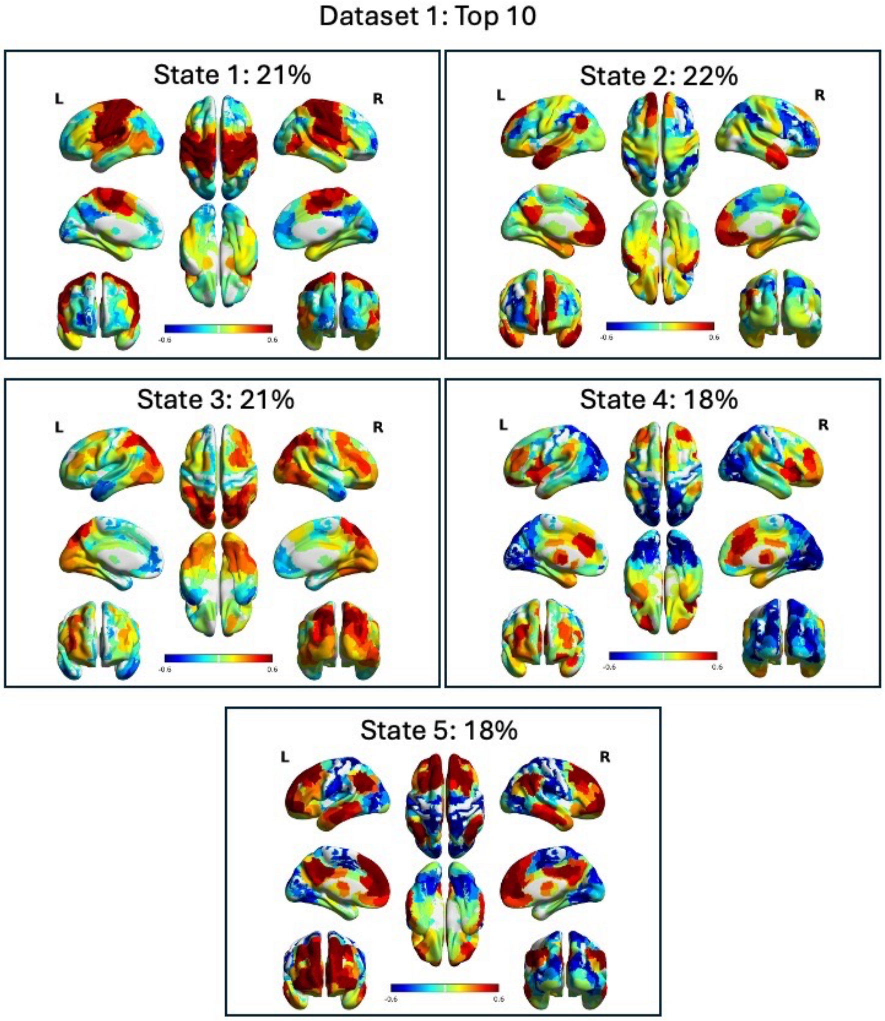

**Supplemental Figure 2**: Dataset 1 dCAPs for low-alpha periods (dLAPs) using a 10% threshold for alpha power. The order of states is arbitrary and does not necessarily match the main manuscript ordering. The frequency that the state occurs is shown in percent.

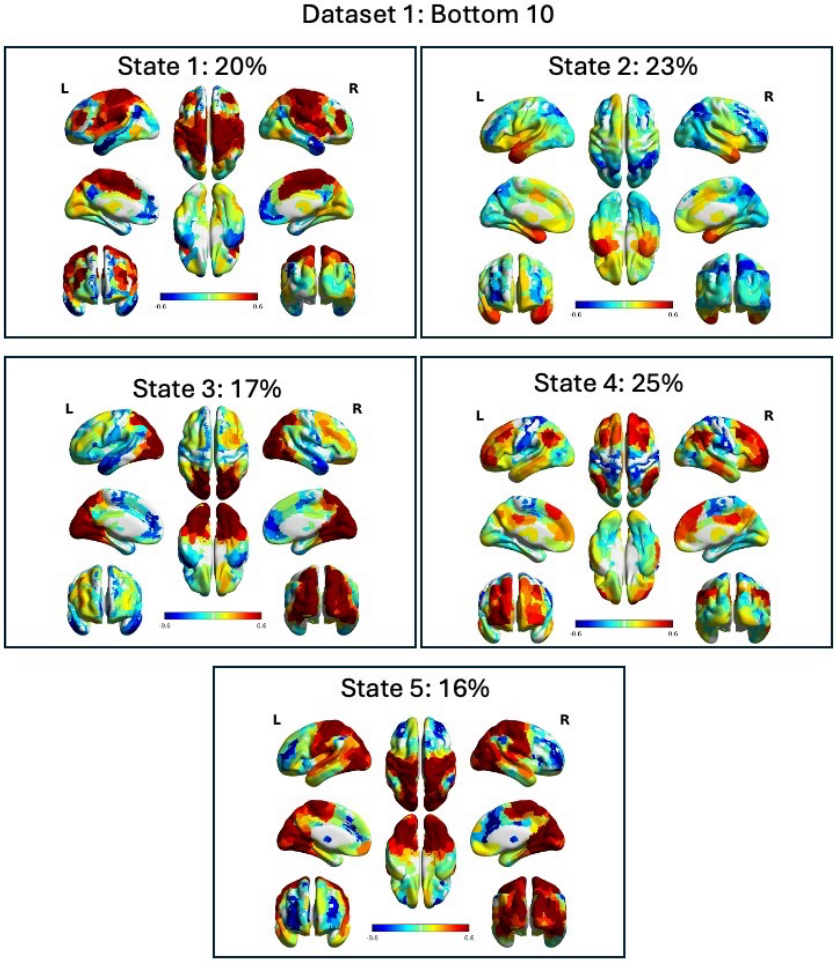

**Supplemental Figure 3**: Dataset 1 dCAPs for high-alpha periods (dHAPs) using a 20% threshold for alpha power. The order of states is arbitrary and does not necessarily match the main manuscript ordering. The frequency that the state occurs is shown in percent.

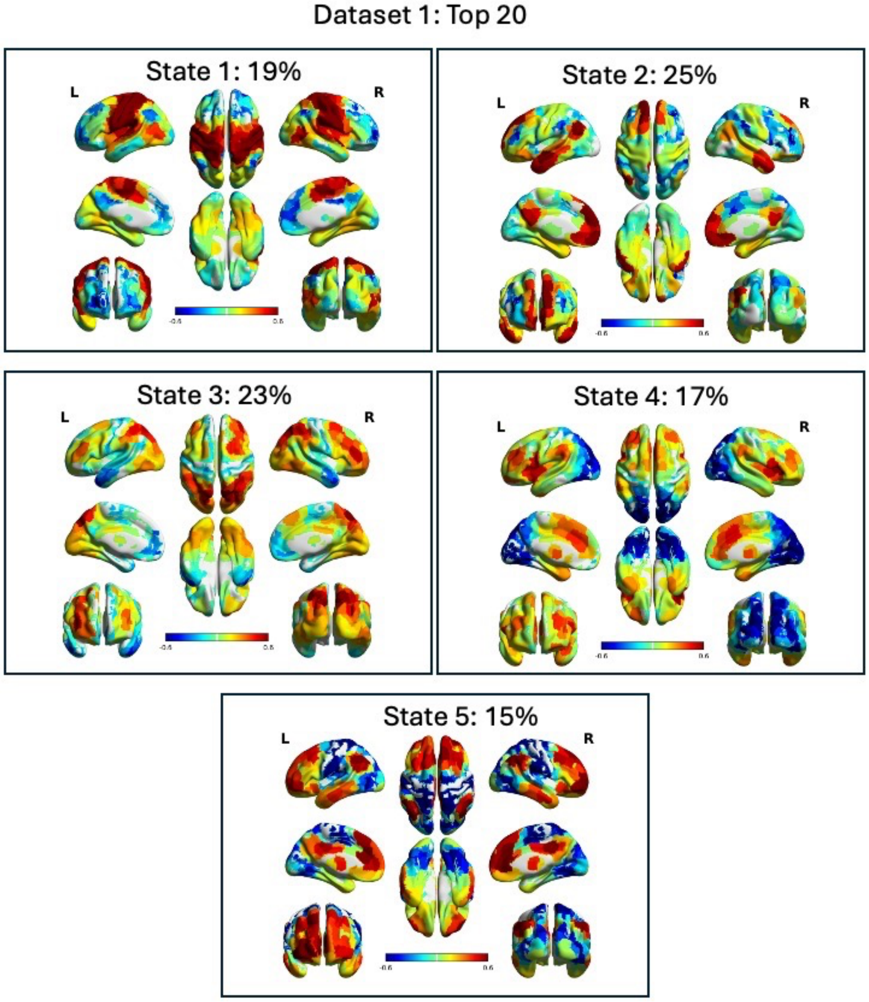

**Supplemental Figure 4**: Dataset 1 dCAPs for low-alpha periods (dLAPs) using a 20% threshold for alpha power. The order of states is arbitrary and does not necessarily match the main manuscript ordering. The frequency that the state occurs is shown in percent.

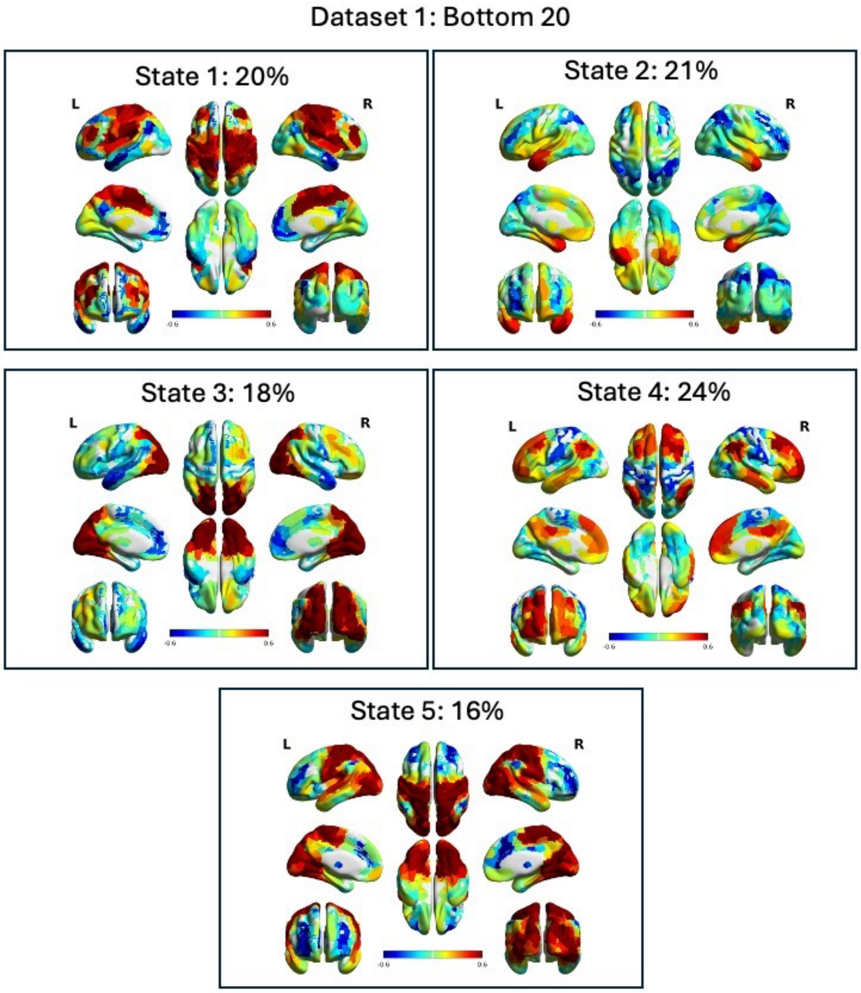

**Supplemental Figure 5**: Dataset 1 K-means clustering of phase-scrambled data using a 15% threshold for high-alpha power (dHAPs). There is no discernable network activity. This demonstrates that non-phase scrambled data is plausibly showing neurological activity.

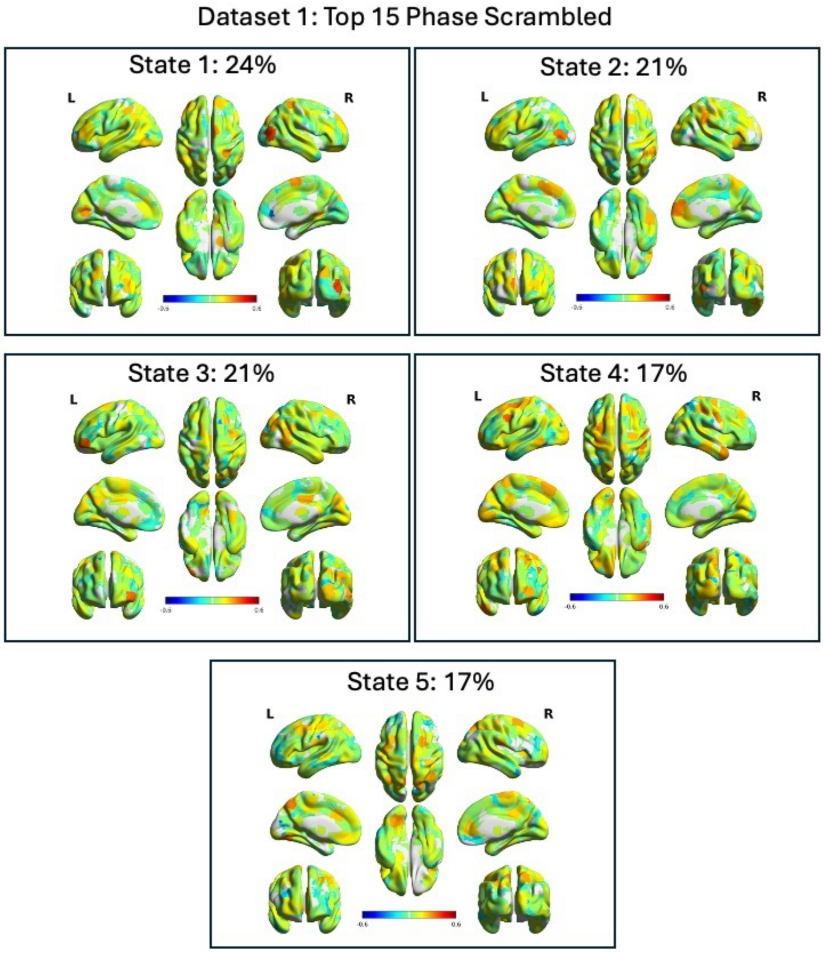

**Supplemental Figure 6**: Dataset 1 K-means clustering of phase-scrambled data using a 15% threshold for low-alpha power (dLAPs). There is no discernable network activity. This demonstrates that non-phase scrambled data is plausibly showing neurological activity.

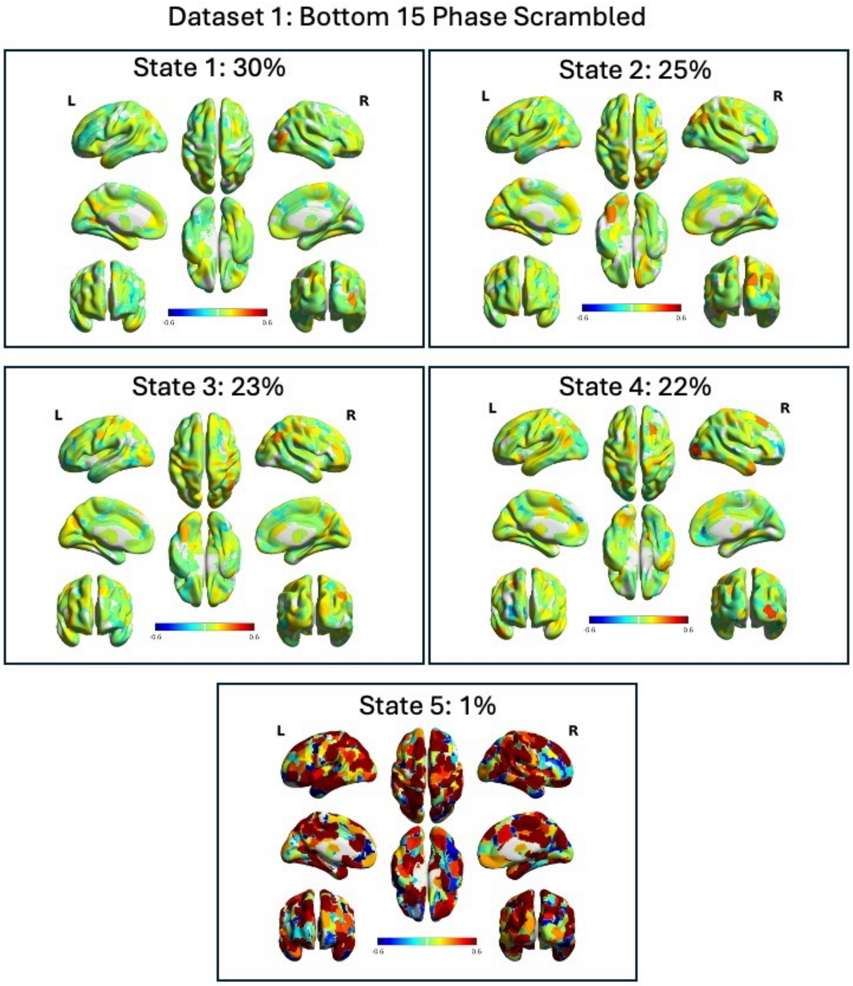

**Supplemental Figure 7:** Dataset 1 K-means clustering of data without global signal regression using a 15% threshold for high-alpha power (dHAPs). Leaving the global signal within the time series produces global states of cortex-wide positive and negative activation.

Dataset 1: Top 15 No GS regression

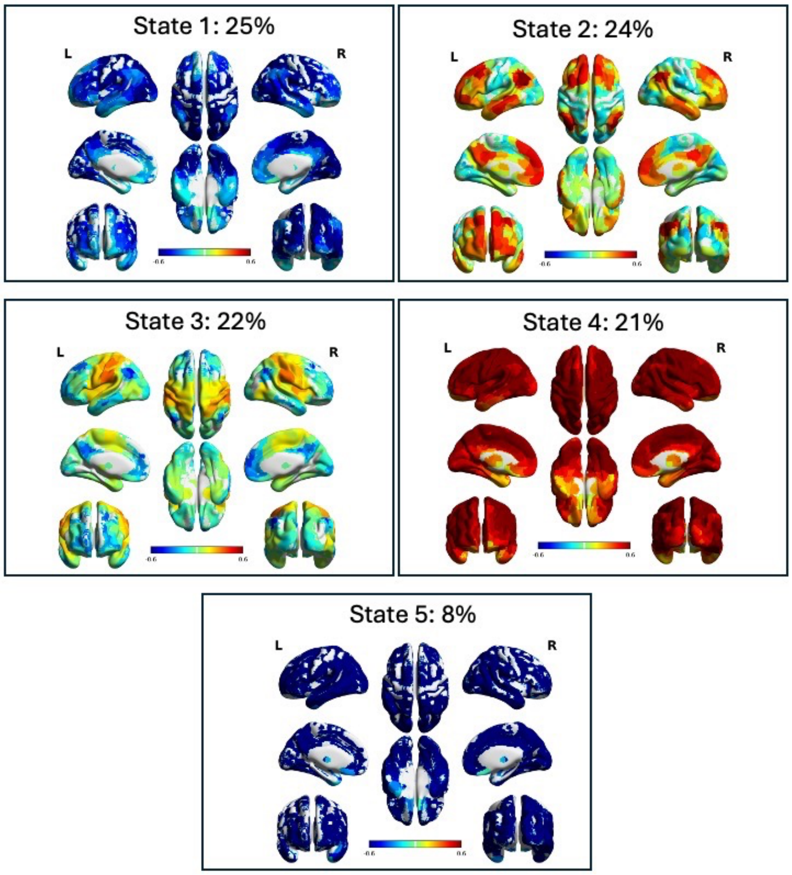

**Supplemental Figure 8:** Dataset 1 K-means clustering of data without global signal regression using a 15% threshold for low-alpha power (dLAPs). Leaving the global signal within the time series produces global states of cortex-wide positive and negative activation.

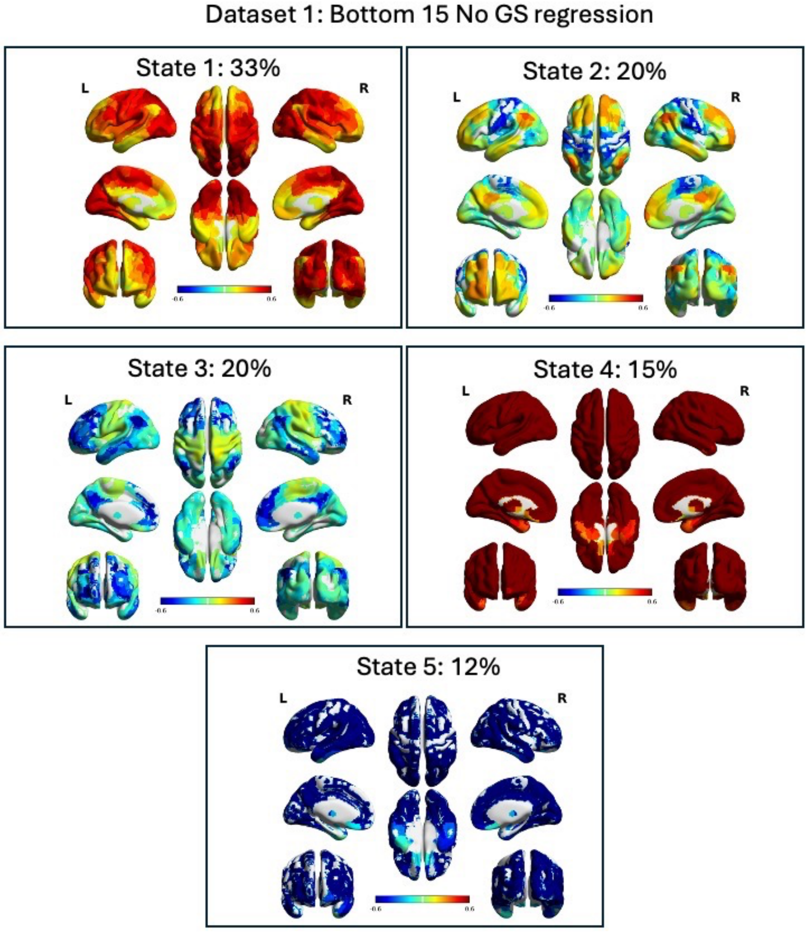

**Supplemental Figure 9**: Dataset 1 k-means clustering of all BOLD TRs using phase-scrambled data. There is no discernable network activity. The order of states is arbitrary and the frequency that the state occurs is shown in percent.

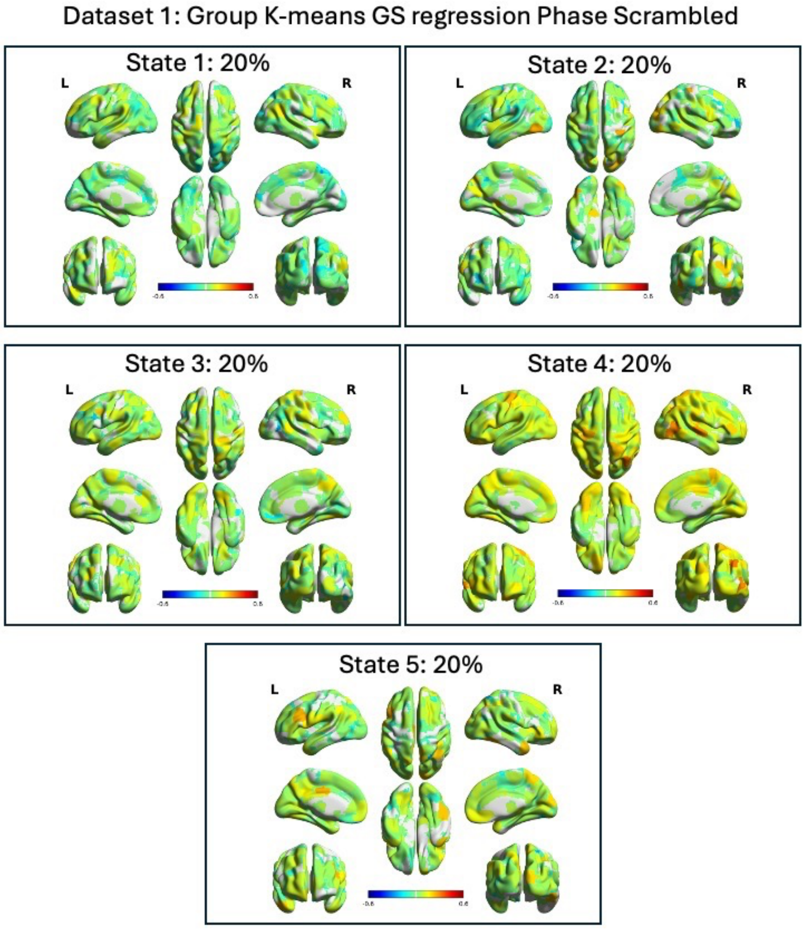

**Supplemental Figure 10:** Dataset 1 group kmeans without global signal regression. Leaving the global signal within the time series produces global states of cortex-wide positive and negative activation.

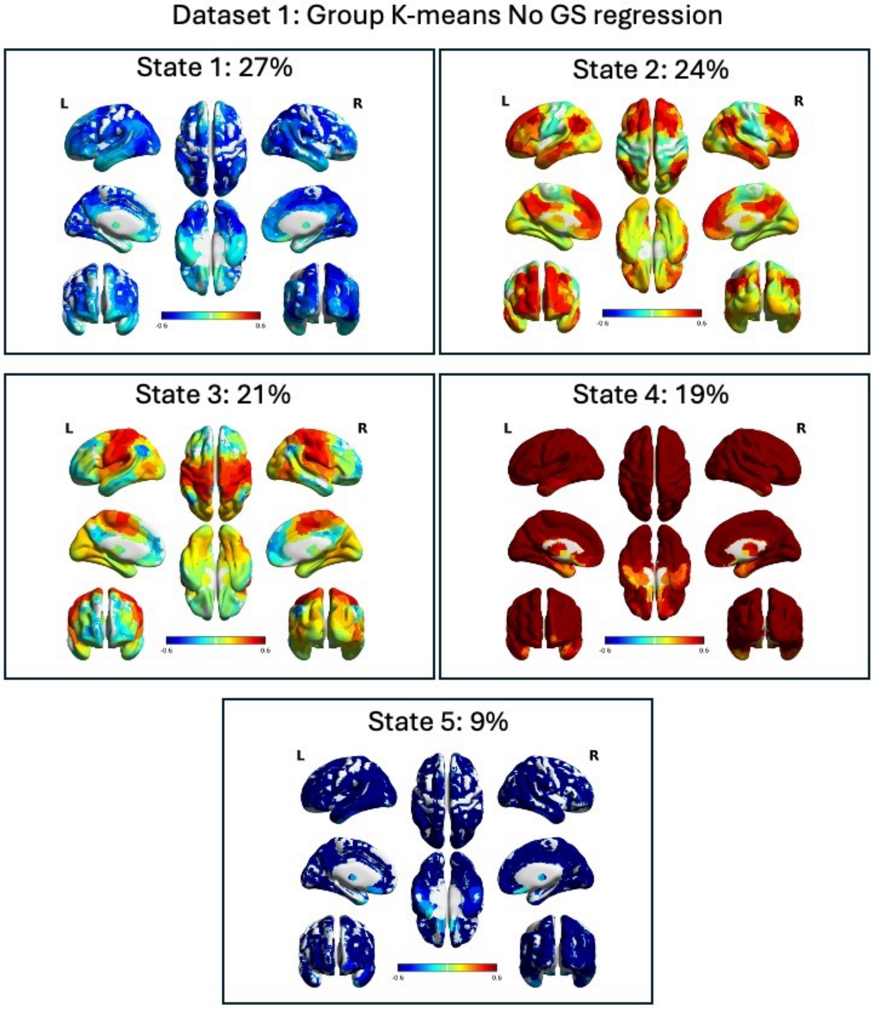

**Supplemental Figure 11:** Dataset 2 k-means clustering of all BOLD TRs using phase-scrambled data. There is no discernable network activity. The order of states is arbitrary and the frequency that the state occurs is shown in percent.

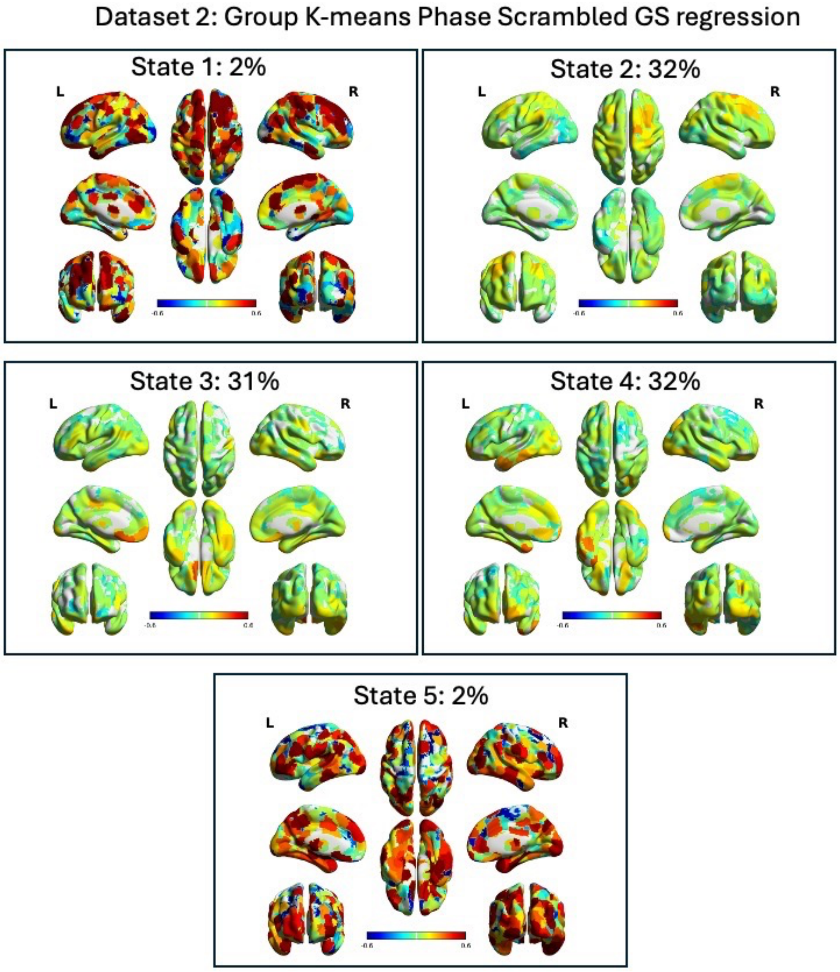

**Supplemental Figure 12:** Dataset 2 group kmeans without global signal regression. Leaving the global signal within the time series produces global states of cortex-wide positive and negative activation.

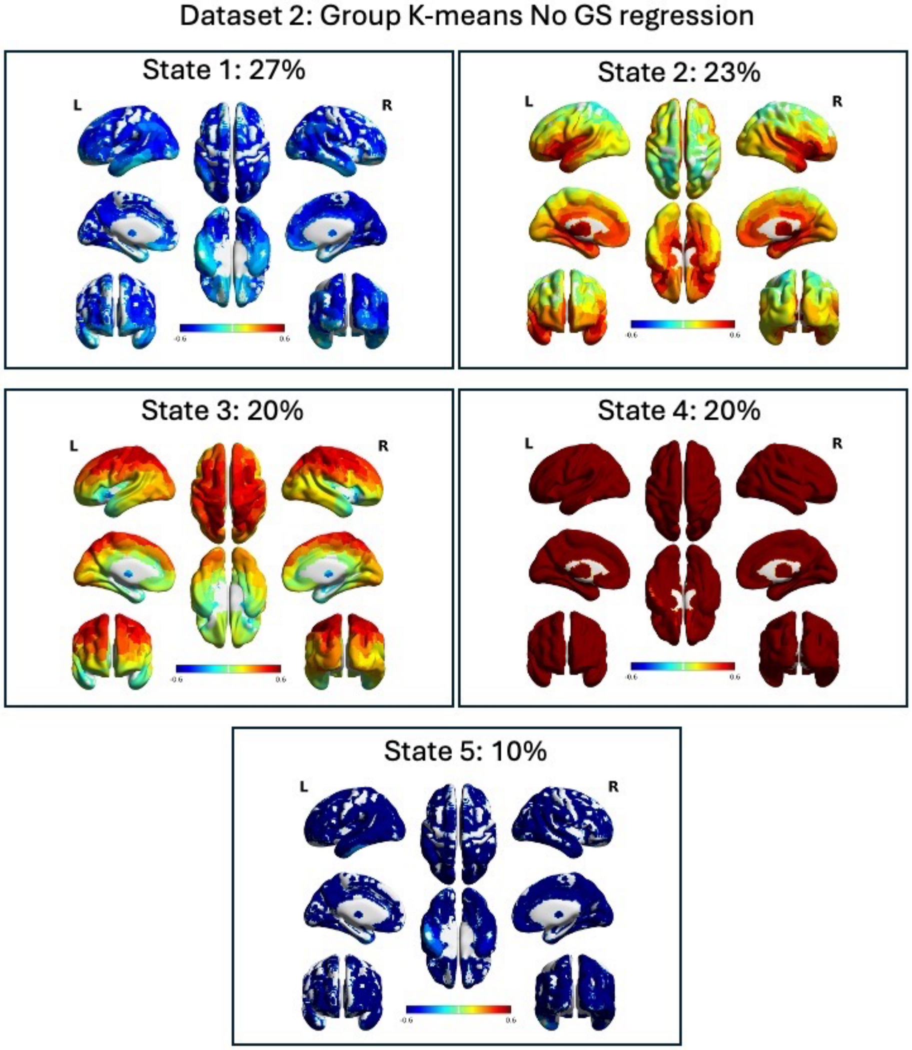

**Supplemental Figure 13:** Dataset 2 dCAPs for high-alpha periods (dHAPs) using a 10% threshold for alpha power. The order of states is arbitrary and does not necessarily match the main manuscript ordering. The frequency that the state occurs is shown in percent.

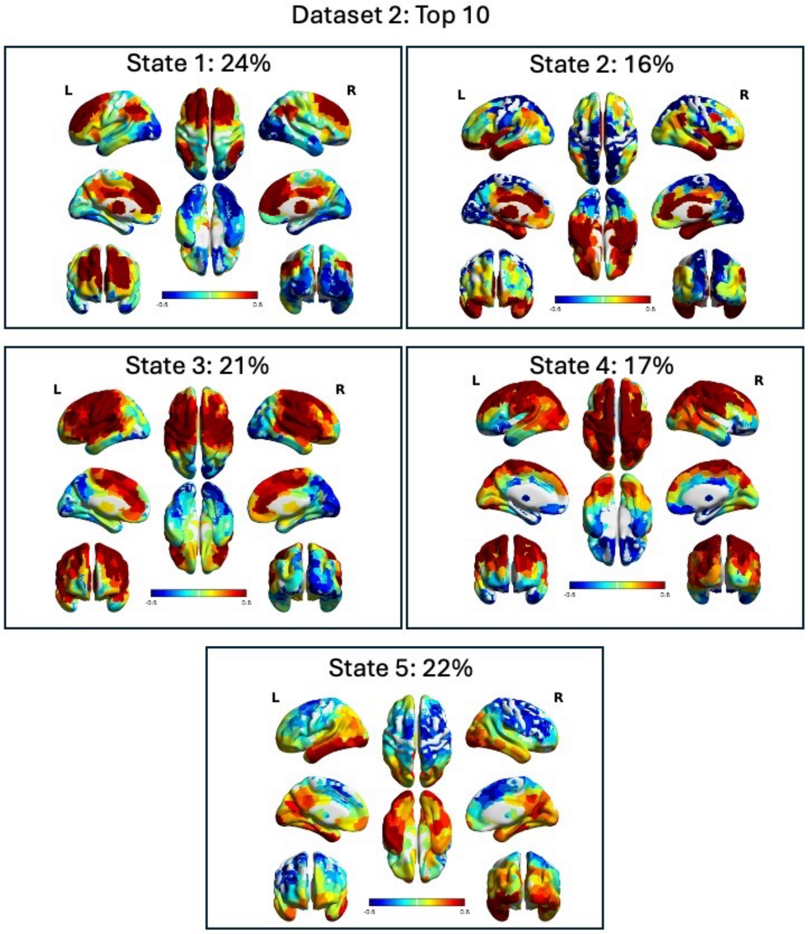

**Supplemental Figure 14:** Dataset 2 dCAPs for low-alpha periods (dLAPs) using a 10% threshold for alpha power. The order of states is arbitrary and does not necessarily match the main manuscript ordering. The frequency that the state occurs is shown in percent.

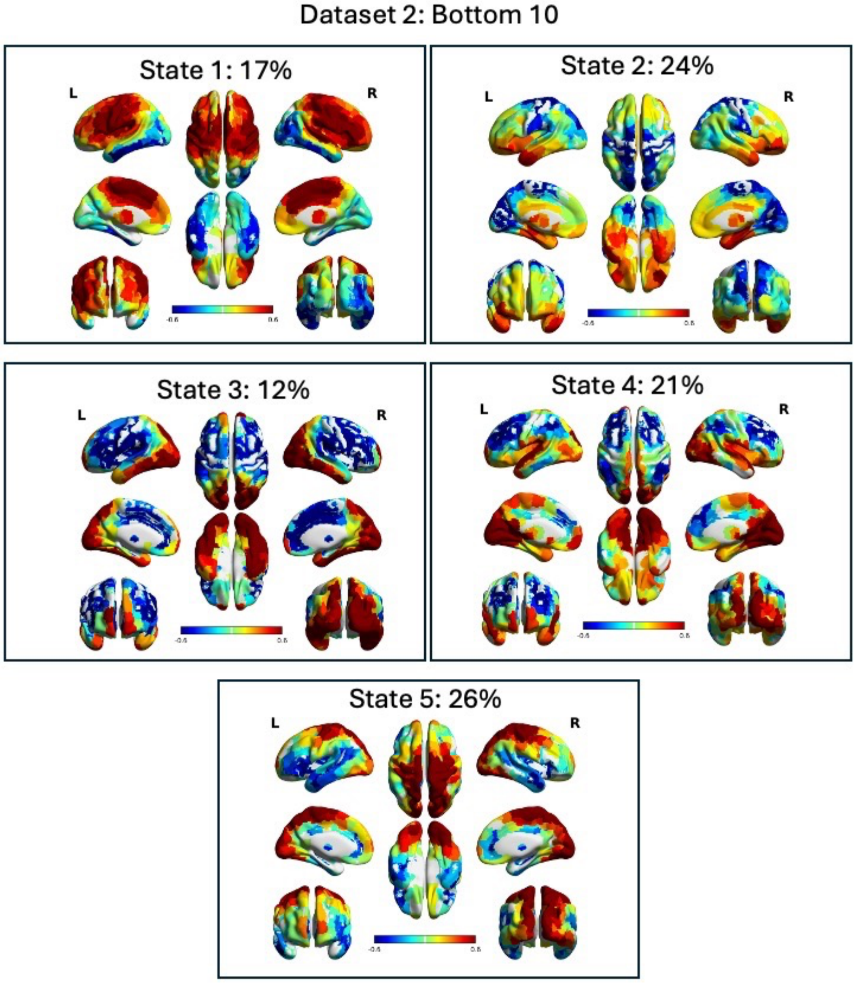

**Supplemental Figure 15**: Dataset 2 dCAPs for high-alpha periods (dHAPs) using a 20% threshold for alpha power. The order of states is arbitrary and does not necessarily match the main manuscript ordering. The frequency that the state occurs is shown in percent.

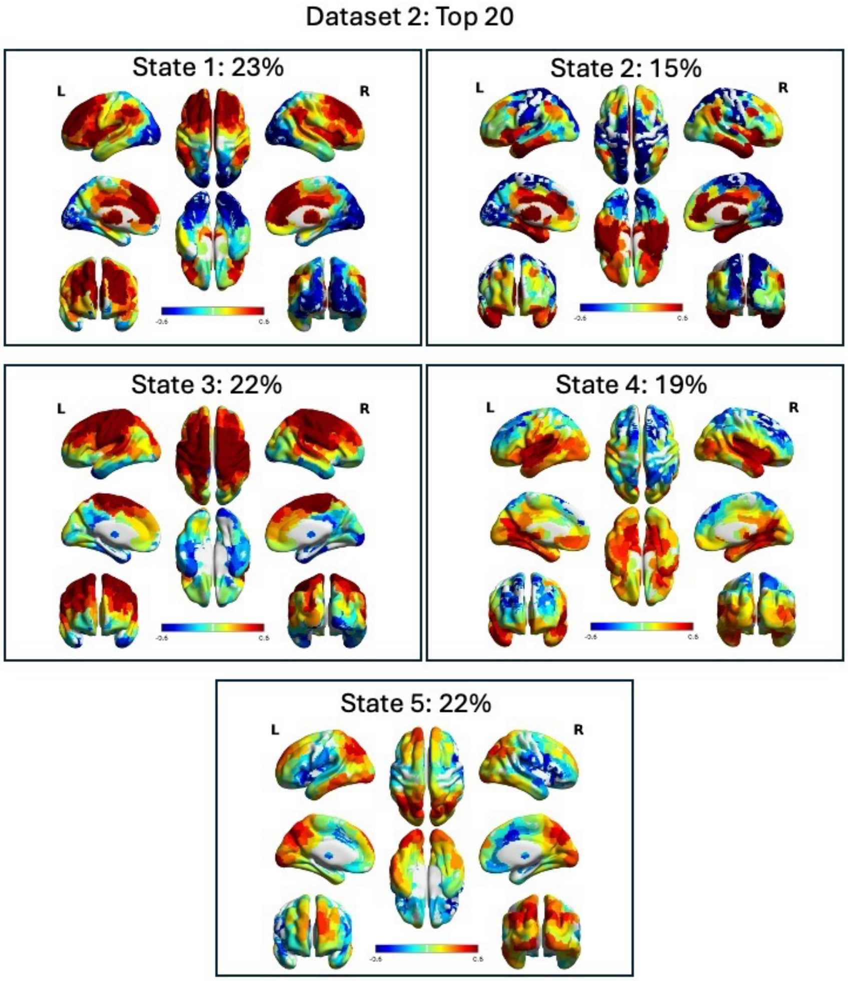

**Supplemental Figure 16**: Dataset 2 dCAPs for low-alpha periods (dLAPs) using a 20% threshold for alpha power. The order of states is arbitrary and does not necessarily match the main manuscript ordering. The frequency that the state occurs is shown in percent.

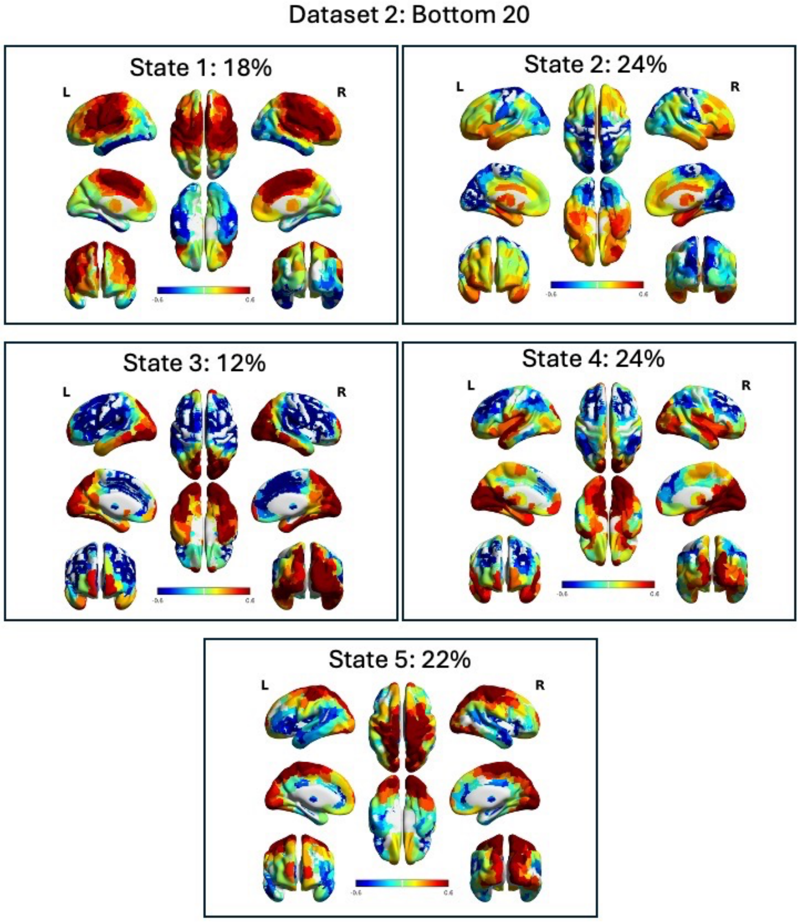

**Supplemental Figure 17**: Dataset 2 K-means clustering of phase-scrambled data using a 15% threshold for high-alpha power (dHAPs). There is no discernable network activity. This demonstrates that non-phase scrambled data is plausibly showing neurological activity.

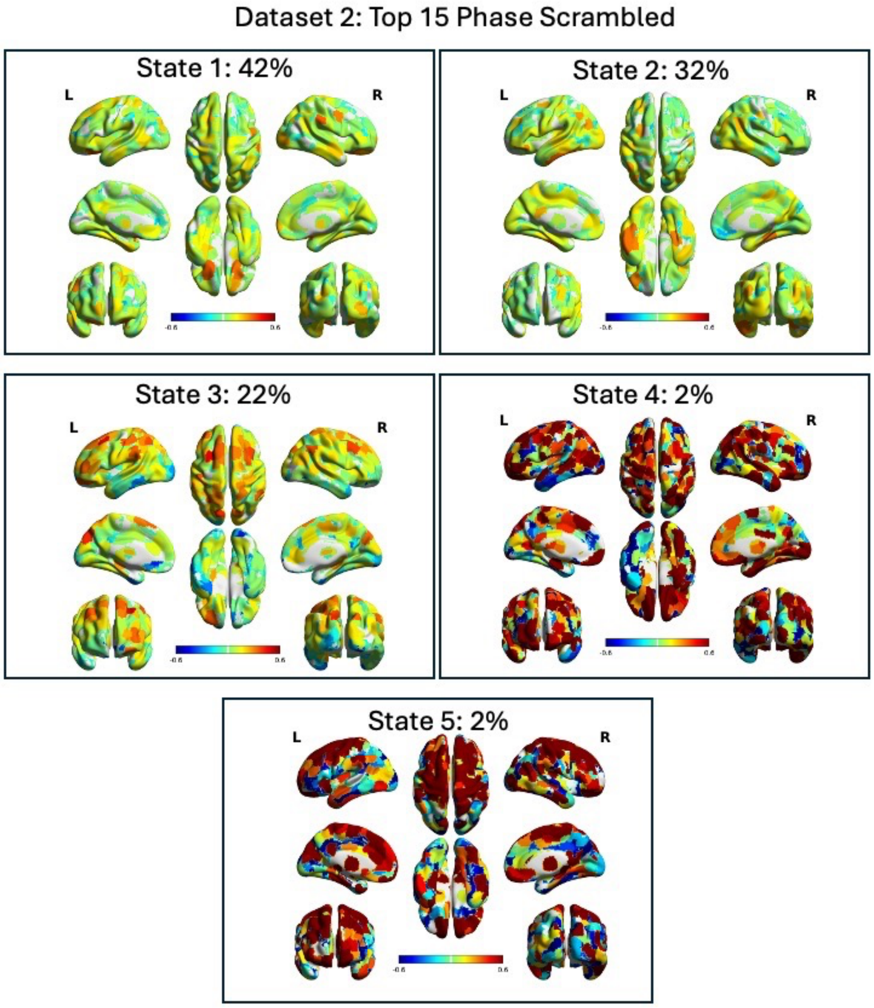

**Supplemental Figure 18**: Dataset 2 K-means clustering of phase-scrambled data using a 15% threshold for low-alpha power (dLAPs). There is no discernable network activity. This demonstrates that non-phase scrambled data is plausibly showing neurological activity.

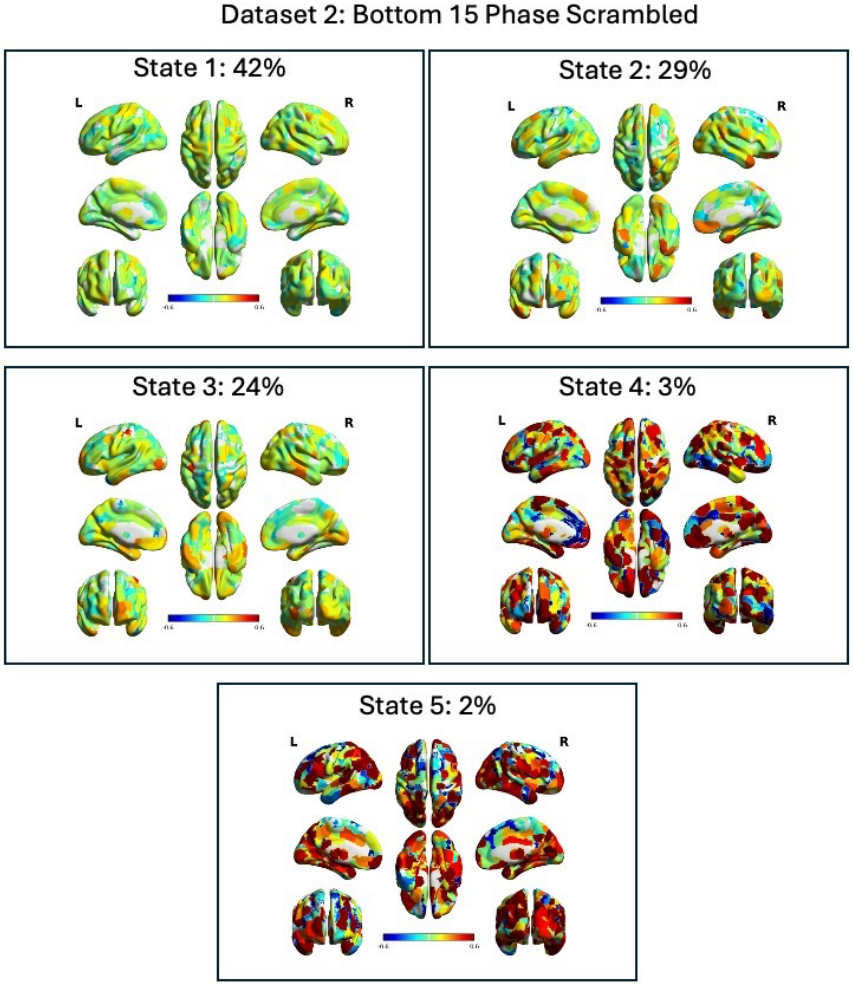

**Supplemental Figure 19:** Dataset 2 K-means clustering of data without global signal regression using a 15% threshold for high-alpha power (dHAPs). Leaving the global signal within the time series produces global states of cortex-wide positive and negative activation.

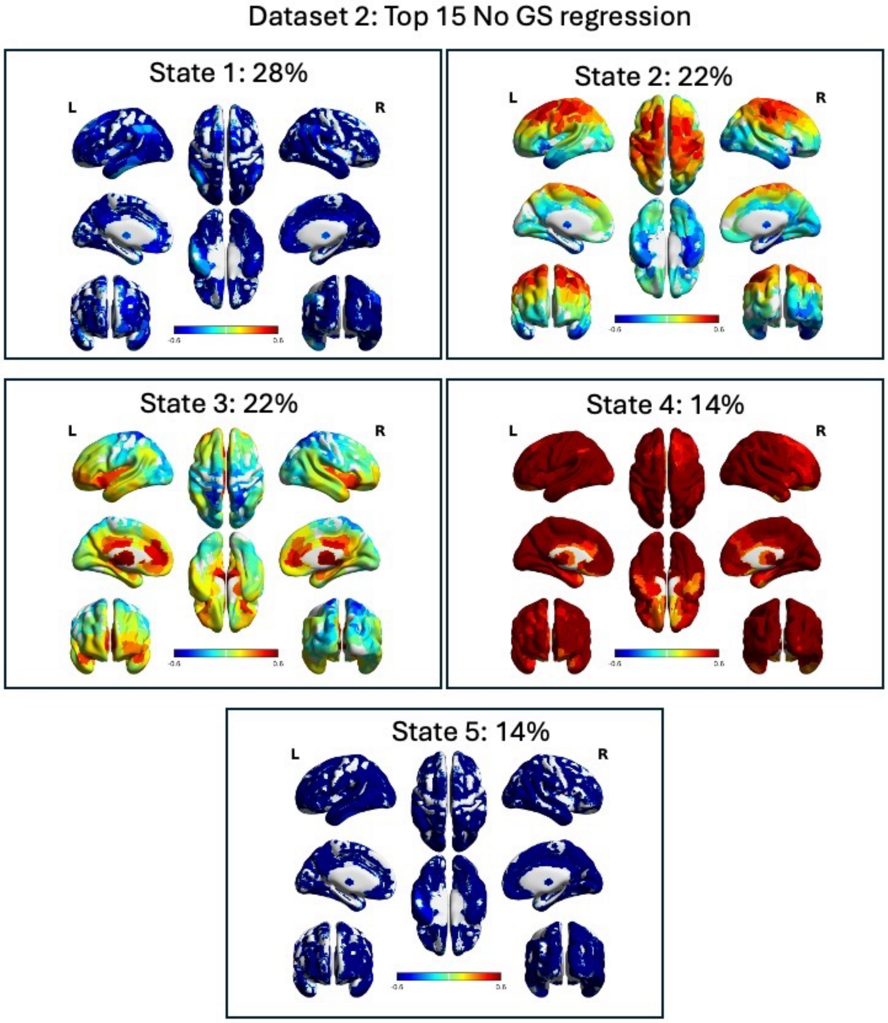

**Supplemental Figure 20:** Dataset 1 K-means clustering of data without global signal regression using a 15% threshold for low-alpha power (dLAPs). Leaving the global signal within the time series produces global states of cortex-wide positive and negative activation.

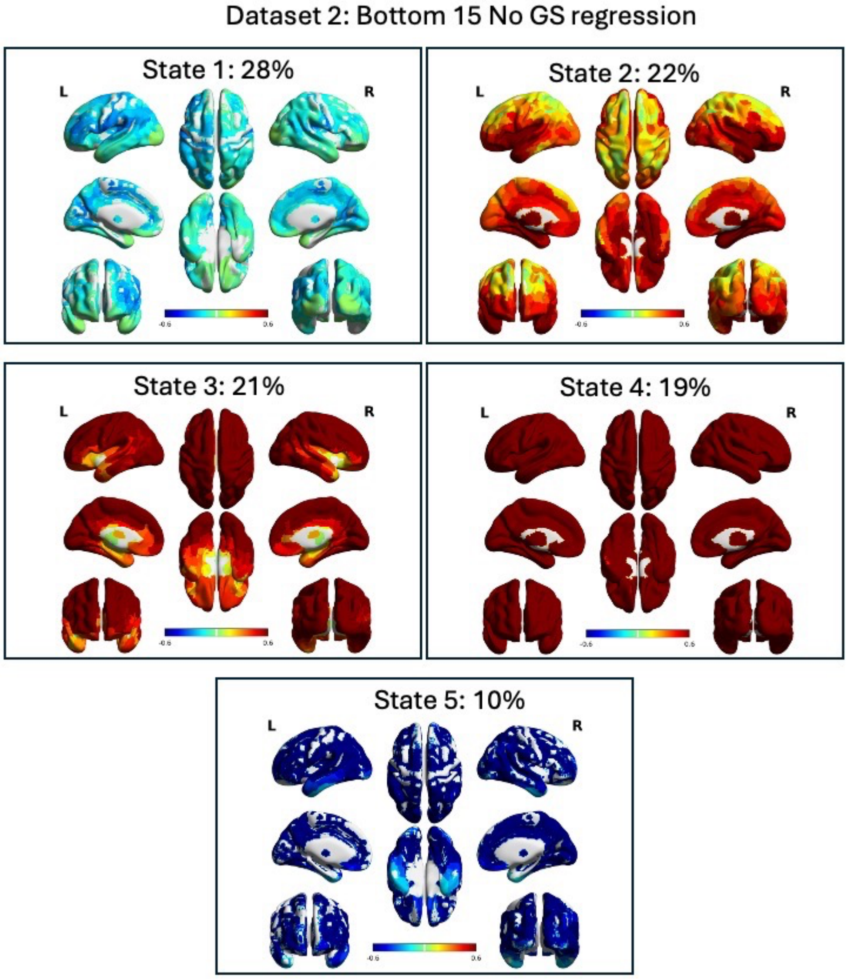

**Supplemental Figure 21:** Static functional connections of the alpha-EEG time series for dataset 1 without global signal regression.**Top**: Unthresholded z-scored spatial map of a one sample t-test on voxel-wise linear regression beta estimates between the alpha-EEG time series and the BOLD signal. Alpha is generally associated negatively with the visual cortex and positively associated with the rest of the cortex. **Bottom**: Cluster corrected results from a GLM with no covariates showed the strongest BOLD associations with alpha are negative and within the visual cortex ($p < 0.001$ voxel-wise uncorrected, $p < 0.05$ cluster-wise corrected).

Dataset 1: Static Functional Connections
of alpha-EEG time-series

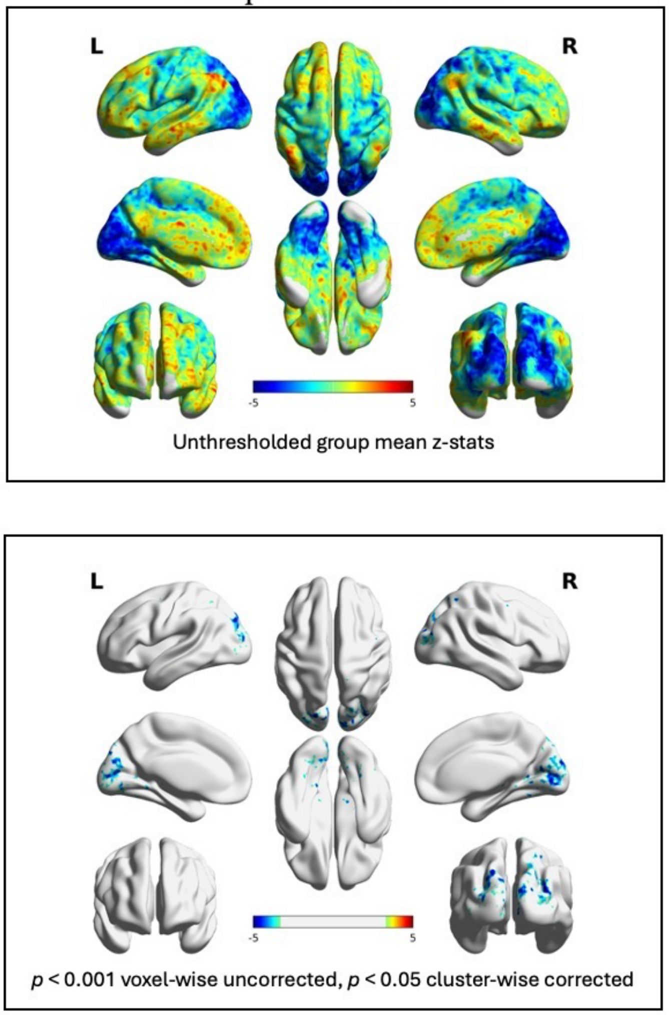

**Supplemental Figure 22:** Static functional connections of the alpha-EEG time series for dataset 1 with global signal regression.**Top**: Unthresholded z-scored spatial map of a one sample t-test on voxel-wise linear regression beta estimates between the alpha-EEG time series and the BOLD signal. Alpha is generally associated negatively with the visual cortex and positively associated with the rest of the cortex. **Bottom**: Cluster corrected results from a GLM with no covariates showed the strongest BOLD associations with alpha are negative and within the visual cortex ($p < 0.001$ voxel-wise uncorrected, $p < 0.05$ cluster-wise corrected).

Dataset 1: Static Functional Connections of alpha-EEG time-series with GS regression

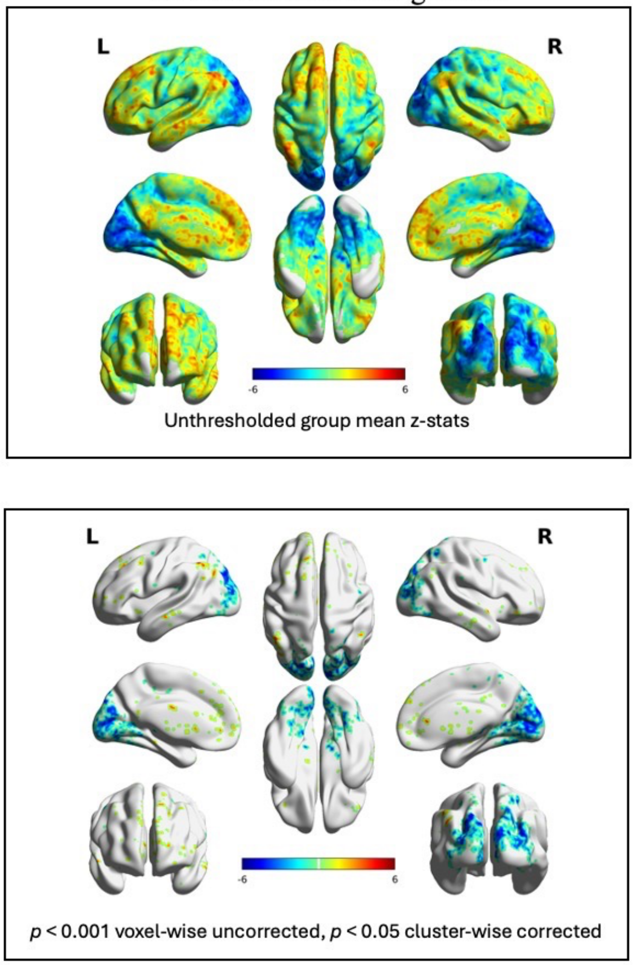

**Supplemental Figure 23:** Static functional connections of the alpha-EEG time series for dataset 2 without global signal regression.**Top**: Unthresholded z-scored spatial map of a one sample t-test on voxel-wise linear regression beta estimates between the alpha-EEG time series and the BOLD signal. Alpha is generally associated negatively with the visual and sensorimotor cortices and positively associated with the rest of the cortex. **Bottom**: Cluster corrected results from a GLM with no covariates showed the strongest BOLD associations with alpha are negative and within the visual cortex ($p < 0.001$ voxel-wise uncorrected, $p < 0.05$ cluster-wise corrected).

Dataset 2: Static Functional Connections of alpha-EEG time-series

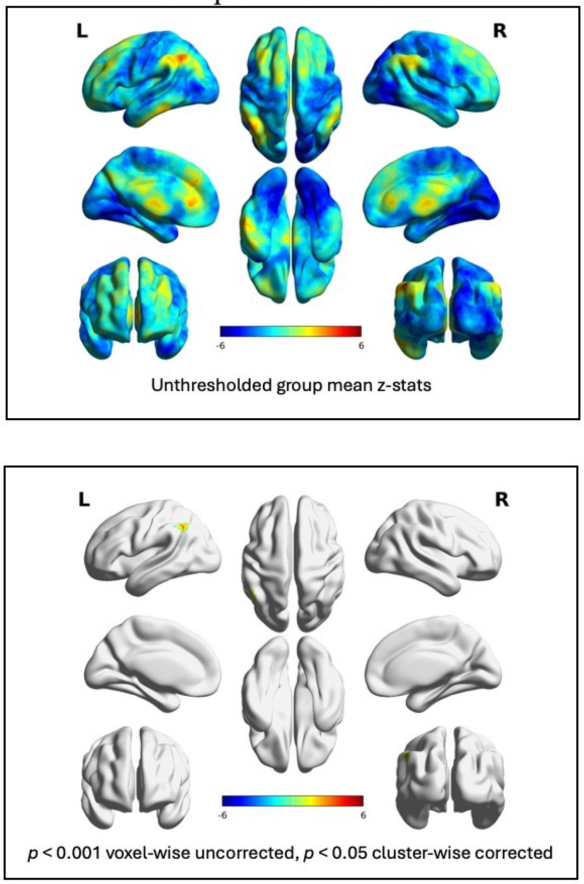

**Supplemental Figure 24:** Static functional connections of the alpha-EEG time series for dataset 2 with global signal regression.**Top**: Unthresholded z-scored spatial map of a one sample t-test on voxel-wise linear regression beta estimates between the alpha-EEG time series and the BOLD signal. Alpha is generally associated negatively with the visual and sensorimotor cortices and positively associated with the rest of the cortex. **Bottom**: Cluster corrected results from a GLM with no covariates showed the strongest BOLD associations with alpha are positive in the parietal cortex, dorsal lateral prefrontal cortex, and the medial prefrontal cortex ($p < 0.001$ voxel-wise uncorrected, $p < 0.05$ cluster-wise corrected).

Dataset 2: Static Functional Connections of alpha-EEG time-series with GS regression

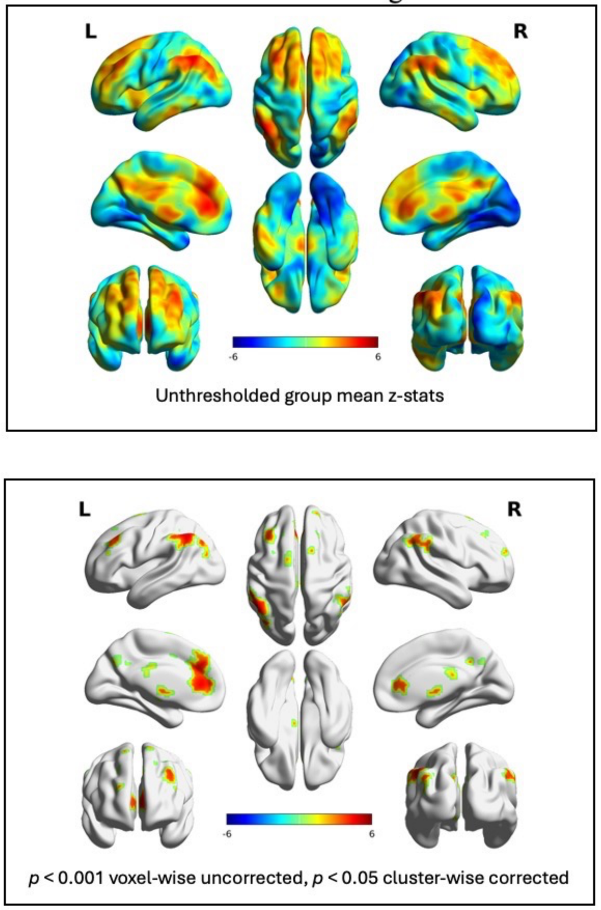

**Supplemental Figure 25:** Static CAP analysis on data with and without global signal regression showing the mean BOLD activation related to the top 15% and bottom 15% of TRs associated with the highest and lowest levels of alpha EEG activity.

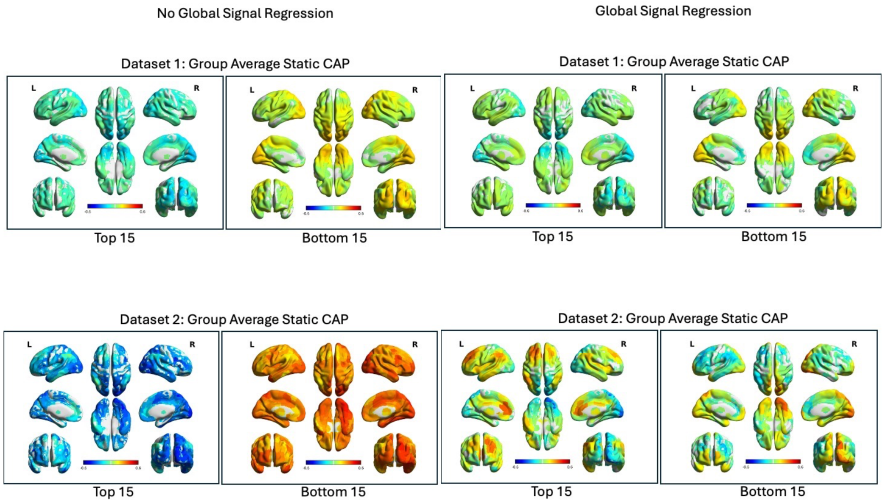

**Supplemental Figure 26:** Elbow criterion for datasets 1 and 2 showing that the optimal cluster solution for all dynamic CAP analyses is 5.

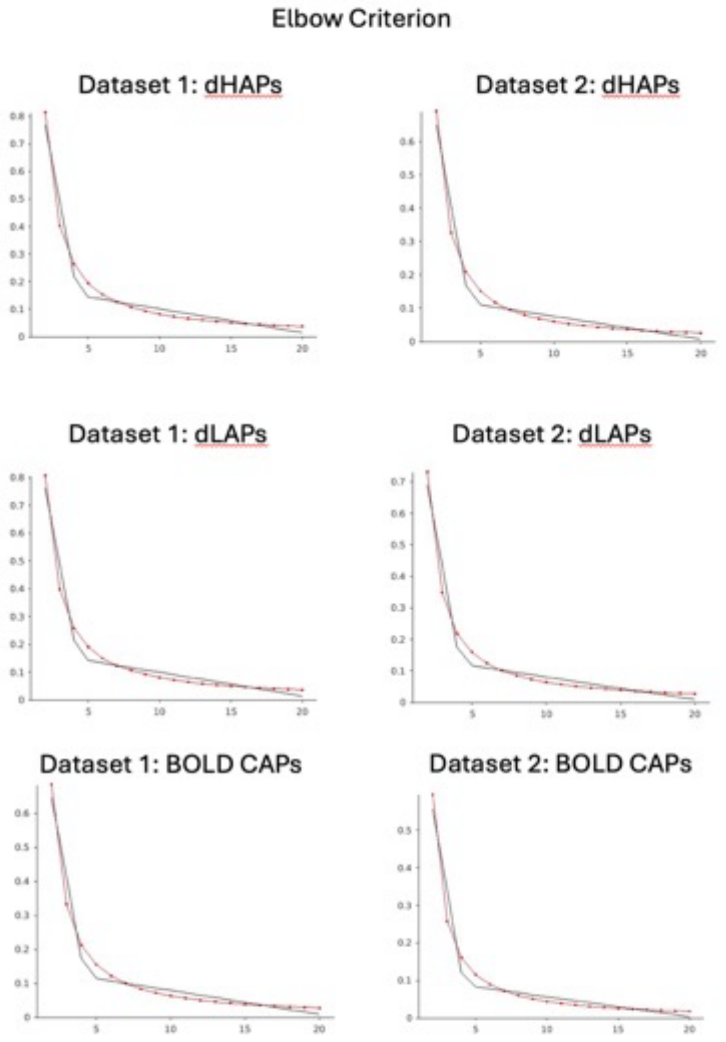
